# PerturbLDM: conditional latent diffusion for modelling single-cell perturbation responses

**DOI:** 10.64898/2026.08.07.743610

**Authors:** Lishan Yu, Kang-Lin Hsieh, Yan Chu, Qizhen Lan, Xingzhong Zhao, Yu-Chun Hsu, Cassidy S. Wood, Laila Rasmy, Patrick G. Pilié, Degui Zhi, Zhongming Zhao, Xiaoqian Jiang, Yulin Dai

**Affiliations:** Center for Secure Artificial Intelligence for Healthcare, Department of Health Data Science and Artificial Intelligence, McWilliams School of Biomedical Informatics, The University of Texas Health Science Center at Houston, Houston, TX 77030, USA; Department of Genitourinary Medical Oncology, Division of Cancer Medicine, UT MD Anderson Cancer Center, Houston, TX 77030, USA; Department of Radiation Physics, Division of Radiation Oncology, UT MD Anderson Cancer Center, Houston, TX 77030, USA; Center for Artificial Intelligence and Genome Informatics, Department of Bioinformatics and Systems Medicine, McWilliams School of Biomedical Informatics, The University of Texas Health Science Center at Houston, Houston, TX 77030, USA; Department of Health Data Science and Artificial Intelligence, McWilliams School of Biomedical Informatics, The University of Texas Health Science Center at Houston, Houston, TX 77030, USA; Center for Precision Health, Department of Bioinformatics and Systems Medicine, McWilliams School of Biomedical Informatics, The University of Texas Health Science Center at Houston, Houston, TX 77030, USA; Department of Biomedical Informatics, Vanderbilt University Medical Center, Nashville, TN 37203, USA

**Author notes:** Correspondence: Yulin Dai, Ph.D., Department of Bioinformatics and Systems Medicine, McWilliams School of Biomedical Informatics, The University of Texas Health Science Center at Houston, 7000 Fannin St., Suite E760A, Houston, TX 77030, USA.

## Abstract

Single-cell perturbation profiling maps intervention-induced phenotypes, yet experiments measure only a fraction of the perturbation-context space. Learning context-dependent perturbation effects could enable response prediction beyond measured conditions. Here we introduce PerturbLDM, a latent-diffusion framework for conditional generation of single-cell transcriptional responses. Following Tahoe-100M pretraining, it predicted 13,942 held-out combinations of observed drugs, doses and cell lines more accurately than existing methods, with higher matched-control effect correlation than an additive marginal baseline in 95.2% of conditions. The Tahoe-100M-pretrained model was further used to rank PANACEA compounds by pathway similarity, placing shared-mechanism pairs among nearest neighbours. In smaller datasets, PerturbLDM generated a mid-gestational fetal-colon state with 67% lower gene-wise error than Squidiff, retaining the balance between absorptive and BEST4/OTOP2-like epithelial programmes. In PBMCs, it captured six of seven interferon and antiviral programmes and the interferon-associated FAO–OXPHOS programme more accurately than scGen. Together, these results support conditional response generation across data scales and biological settings.

## Main

Perturbational transcriptomic atlases enable systematic, large-scale characterisation of how molecular and biological interventions reshape cellular states. Bulk resources such as the Connectivity Map and LINCS established that gene-expression signatures can connect compounds, mechanisms and transcriptional phenotypes^1,2^. Single-cell screens extend this approach by resolving perturbation-specific gene programmes, heterogeneous responses and context-dependent cellular states ^3–9^. Despite advances in profiling scale and resolution, comprehensive measurement across all combinations of perturbation, dose and cellular context remains experimentally infeasible ^9–11^. Computational models could therefore inform experimental design by estimating cellular responses for conditions not yet profiled. Such use depends on recovering intervention-induced transcriptional effects across cellular contexts rather than cellular background alone ^12,13^. This places perturbation-response prediction among the central objectives of virtual-cell modelling ^14^. Recent benchmarks emphasise marginal baselines and direct effect metrics because absolute-expression metrics alone can obscure response fidelity^12,13^.

Various methods have been developed for perturbation-response prediction. Variational autoencoder (VAE)-based models such as scGen learn low-dimensional representations of cellular states and use latent-space vector arithmetic to transfer perturbation effects across cellular contexts ^15^. CPA and chemCPA instead separate basal state from perturbation and covariate representations and use dose-dependent scaling, allowing these components to be recombined for unmeasured condition combinations. ChemCPA additionally derives drug representations from molecular structure ^10,16^. Unpaired cell populations can be modelled with optimal transport, whereas graph-based methods use gene relationships to predict combinatorial genetic perturbations ^11,17^. In parallel, foundation models such as Geneformer, scGPT and scFoundation use large-scale Transformer pretraining to learn transferable representations for downstream single-cell tasks ^18–20^. Although these models show broad potential, their pretraining objectives are not necessarily optimised for direct prediction of continuous, condition-specific gene-expression values. More recently, diffusion models such as Squidiff have used iterative denoising to generate continuous, high-dimensional cellular expression states^21^. Conditional generation of single-cell transcriptional responses for unmeasured conditions has not been established across both atlas-scale perturbation resources and biologically distinct study-specific datasets^9,13^.

Here we introduce PerturbLDM, a conditional latent-diffusion framework for generating single-cell transcriptional responses under specified perturbation and context variables^22^. We evaluated whether its predictions recovered intervention-induced gene effects and coordinated transcriptional programmes rather than absolute expression alone. Condition-aligned local cellular distributions provided a complementary assessment of response fidelity.

We used Tahoe-100M for pretraining and systematic evaluation of conditional generation on held-out combinations of observed factors. In PANACEA, response profiles from a Tahoe-pretrained model variant were used to characterise transcriptional programmes and organise pharmacological relationships^23^. Separately, we fitted task-specific PerturbLDM models to fetal-colon and PBMC datasets to evaluate conditional generation at the scale and design of individual biological studies^24,25^.

## Results

### Conditional latent denoising generates transcriptomic responses

PerturbLDM combines a variational autoencoder with conditional latent diffusion (Fig. 1a). The encoder maps measured single-cell expression profiles to a compact latent space. The diffusion model samples response states through condition-guided denoising, and the decoder returns these states to gene-expression space. Conditioning variables vary across datasets, but each task generates condition-specific single-cell profiles for a response state withheld from model fitting.

**Fig. 1.**
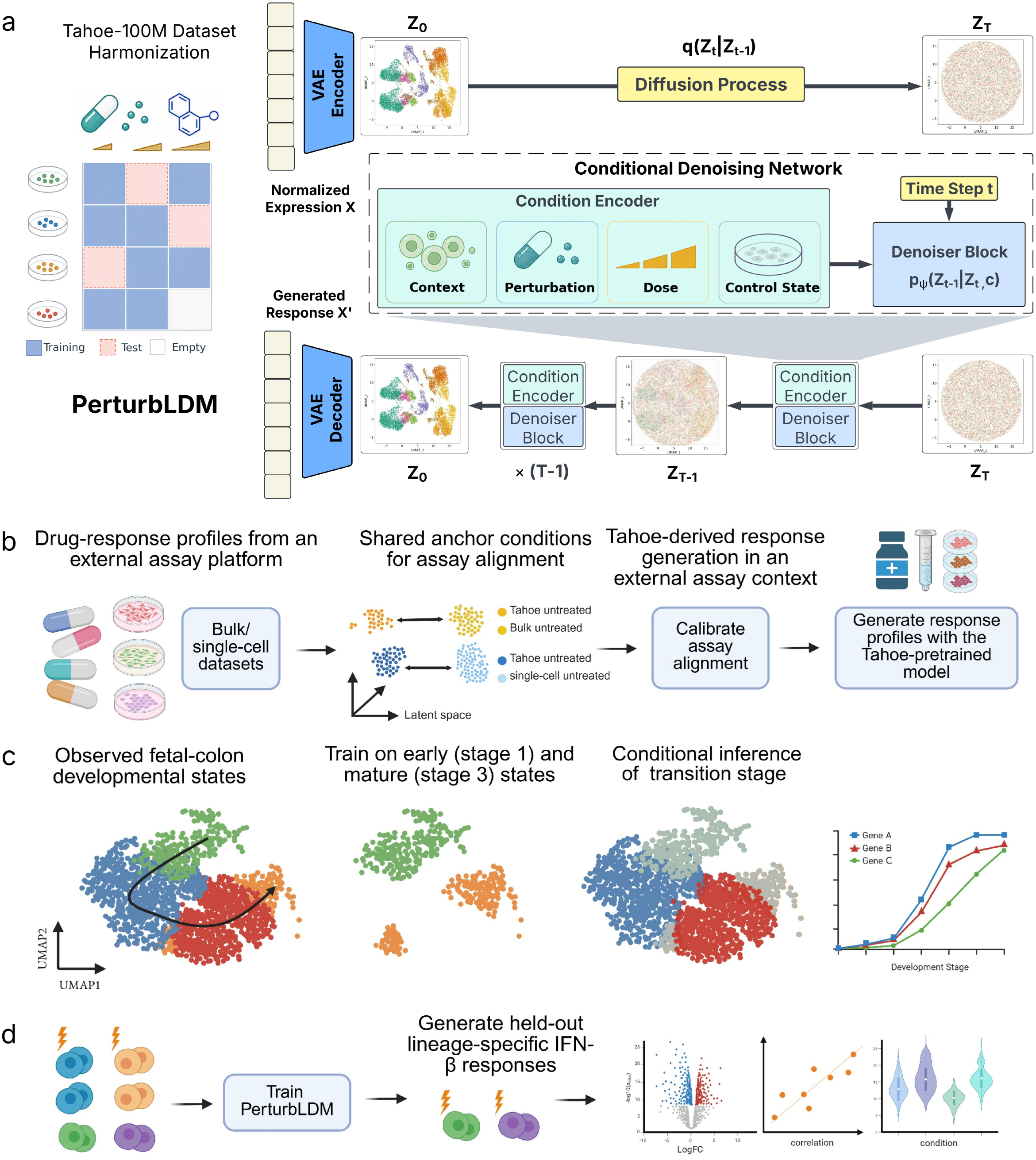
— PerturbLDM framework and scale-to-application evaluation design. a, Tahoe-100M provides the data scale and condition coverage for model training and held-out evaluation. A variational autoencoder encodes measured single-cell profiles, and conditional latent diffusion denoises latent states under context, perturbation, dose and control-state conditioning before decoding the requested response. **b,** PANACEA illustrates a pretrained-model application in which Tahoe-derived response profiles are calibrated to an external assay and used to organise compounds by shared pathway effects. **c,** The fetal-colon application uses early (broad stage 1) and mature (broad stage 3) enterocytes as developmental anchors and evaluates generation of one pooled held-out mid-gestational population (broad stage 2), without PCW-specific conditioning. **d,** The PBMC application evaluates generation of three held-out lineage-specific IFN-*β* responses from retained lineage-matched unstimulated controls.

The evaluation design separated downstream reuse of a pretrained response model from task-specific fitting. Tahoe-100M supported model fitting and the central held-out benchmark. Precomputed response profiles from a Tahoe-trained model variant were subsequently analysed in PANACEA (Fig. 1a,b). In smaller study-specific settings, separately fitted models generated a withheld intermediate developmental state from early and mature anchors, and unmeasured IFN-*β*-stimulated states from retained unstimulated counterparts (Fig. 1c,d).

### Benchmarking held-out drug-response prediction in Tahoe-100M

Reliable conditional generation could provide testable response hypotheses for unmeasured perturbation contexts and help prioritise direct experimental follow-up. We therefore used Tahoe-100M to quantify the accuracy of PerturbLDM predictions for drug-dose-cell-line conditions withheld from training ^9^. Of 46,471 measured conditions, 32,529 were used for training and 13,942 were randomly withheld for evaluation (∼30%, Fig. 2a and Supplementary Fig. S3a–d).

**Fig. 2.**
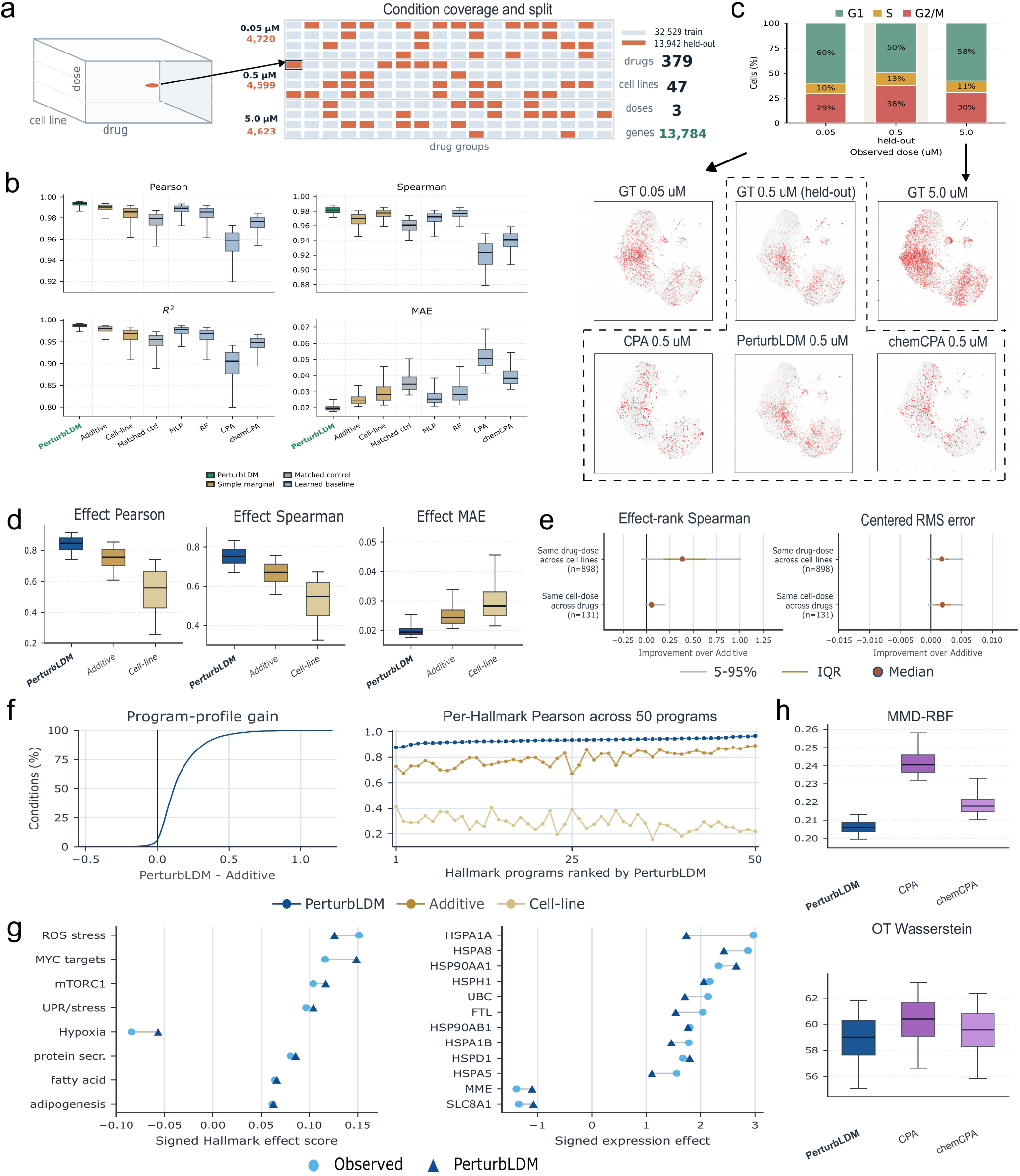
— Benchmarking conditional response generation in Tahoe-100M. a, Retrospective Tahoe benchmark. Approximately 30% of measured treatment conditions (13,942) were randomly withheld, and 32,529 conditions were used for training. All 379 drugs, 47 cell lines and three dose levels occurred in both splits, so the held-out targets comprised unmeasured combinations of observed factors. **b,** Condition-level absolute-expression metrics across the held-out conditions. PerturbLDM is compared with matched control, train-set marginal controls, MLP, random forest (RF), CPA and chemCPA using Pearson correlation, Spearman correlation, *R*^2^ and MAE. Learned-comparator selection used training-derived condition-level validation data and did not use the held-out responses. **c,** Selected Goserelin response in HCT15 cells. Cell-cycle redistribution at the held-out 0.5 *µ*M dose is shown relative to the measured 0.05 and 5.0 *µ*M endpoints, together with observed and predicted state maps. **d,** Treatment-effect recovery after matched-control subtraction, assessed by all-gene Pearson correlation, Spearman correlation and MAE. **e,** Context-specific recovery beyond AdditiveMean. Within each fixed drug-dose or cell-line-dose stratum, dots show the median condition-level gain, thick intervals the interquartile range and thin intervals the 5th–95th percentiles. Effect-rank Spearman gains are PerturbLDM minus AdditiveMean and centred-RMSE reductions are AdditiveMean minus PerturbLDM, so positive values favour PerturbLDM for both metrics. **f,** Recovery of the direction and magnitude of coordinated transcriptional responses across 50 signed Hallmark programmes, shown as condition-level gains over AdditiveMean and per-programme correlations. **g,** Observed and PerturbLDM-predicted signed effects for selected Hallmark programmes and response genes in one illustrative Bortezomib-treated LOX-IMVI condition at 0.05 *µ*M. This illustrative condition was not used for mechanistic inference. **h,** Condition-aligned MMD-RBF and OT Wasserstein distances for local empirical cell-state distributions generated by PerturbLDM, CPA and chemCPA. Lower values indicate closer local alignment, not global calibration of the full single-cell distribution. Boxes in **b**, **d** and **h** show interquartile ranges, centre lines show medians and whiskers span the 5th–95th percentiles. For effect, signed-Hallmark and distribution comparisons, the held-out condition is the paired unit (*n* = 13,942), with Goserelin and Bortezomib shown as selected illustrations. Complete paired estimates, confidence intervals and multiplicity-adjusted tests are reported in Supplementary Table S4.

All 379 drugs, 47 cell lines and three dose levels occurred in both splits. This split evaluated held-out combinations of observed factors rather than extrapolation to unseen drugs, cell lines or dose levels. For each held-out condition, PerturbLDM received the requested drug identity and dose together with the matched DMSO control profile for the target cell line, but not the measured perturbed-cell profiles.

Absolute-profile concordance reflects both stable cell-line identity and marginal drug-dose structure. We compared PerturbLDM with training-set marginal baselines and learned models to quantify condition-specific response recovery beyond these components. AdditiveMean combines training-set cell-line and drug-dose marginal means with a global-mean correction, providing a direct test of gains beyond marginal additivity^12,13^. MLP, random forest, CPA and chemCPA were included as learned comparators^10,16^. Their model settings were selected using a shared condition-level validation subset of the training data (Supplementary Fig. S3e–h).

Across 13,942 held-out conditions, PerturbLDM reproduced the measured expression profiles more accurately than the other methods on each metric in Fig. 2b. AdditiveMean yielded a mean *R*^2^ of 0.9761, Pearson correlation of 0.9888 and Spearman correlation of 0.9668, indicating that cell-line background and train-set drug-dose marginals captured much of the absolute-profile variation. Relative to this baseline, PerturbLDM increased mean *R*^2^ to 0.9853 and Spearman correlation to 0.9807, while reducing mean MAE from 0.0256 to 0.0203, a 20.7% relative reduction. Pearson correlation increased more modestly, from the already near-ceiling value of 0.9888 to 0.9928. Among the learned comparators, MLP had the highest mean *R*^2^, Pearson correlation and lowest MAE (0.9713, 0.9866 and 0.0272), whereas random forest had the highest Spearman correlation (0.9744). PerturbLDM exceeded these values on all four metrics (mean *R*^2^ = 0.9853, Pearson *r* = 0.9928, Spearman *ρ* = 0.9807 and MAE= 0.0203), including a 25.5% lower MAE than MLP. CPA and chemCPA yielded mean *R*^2^ values of 0.8919 and 0.9396, Pearson correlations of 0.9537 and 0.9723, and MAEs of 0.0524 and 0.0402, respectively. The near-ceiling correlations of the marginal baseline underscored the limited specificity of absolute-expression metrics for treatment-effect recovery.

To distinguish intervention-specific recovery from absolute-profile concordance, we centred observed and predicted profiles on their matched DMSO controls. The resulting effect profiles quantified gene-level responses specific to each drug-dose-cell-line context. AdditiveMean remained the reference because it does not model the context-specific interaction between its training-set marginals. Across the 13,942 paired held-out conditions, the matched-control-relative gene effects predicted by PerturbLDM had higher linear concordance in 95.23% of conditions, higher rank concordance in 98.01% and lower effect-magnitude error in 98.98% (Fig. 2d). The median within-condition gains were 0.08085 for effect Pearson correlation (95% bootstrap CI, 0.07957–0.08215) and 0.07903 for effect Spearman correlation (0.07804–0.08000), while effect MAE decreased by 0.004670 (0.004617–0.004725). All three comparisons remained significant after BH correction (two-sided paired Wilcoxon, *P <* 10*^−^*^300^). To determine whether these gains were concentrated in a narrow subset of perturbations or cellular contexts, we also compared the methods within 898 predefined drug-dose groups and 131 predefined cell-line-dose groups. PerturbLDM had lower centred RMSE in 91.5% and higher effect-rank Spearman correlation in 92.4% of drug-dose groups. The corresponding fractions across cell-line-dose groups were 91.6% and 92.4% (Fig. 2e and Supplementary Fig. S5). These gains were observed across both drug-dose and cell-line-dose strata. These summaries quantify variation across held-out conditions for a single fitted model, rather than run-to-run or atlas-to-atlas variability.

Perturbation effects at individual genes can be subtle, whereas aggregating them across functionally related genes can reveal coordinated response programmes. We summarised each condition using signed scores for 50 predefined Hallmark gene sets (Fig. 2f). For each set, the score was the mean matched-control-relative expression effect across overlapping member genes. Its sign indicated whether the genes shifted upward or downward on average, and its magnitude represented the average shift. Relative to AdditiveMean, Hallmark-profile Pearson correlation was higher for PerturbLDM in 94.36% of the 13,942 paired conditions, and Hallmark-score MAE was lower in 87.05%. The median within-condition Pearson gain was 0.1162 (95% bootstrap CI, 0.1143–0.1187), while the median MAE reduction was 0.003640 (0.003555–0.003707). Both paired Wilcoxon comparisons yielded BH-adjusted *P <* 10*^−^*^300^. The improvement extended from individual gene effects to the direction and magnitude of coordinated gene-programme responses. In a selected Bortezomib condition at 0.05 *µ*M in LOX-IMVI cells, the predicted scores followed the observed directions for the displayed ROS stress, MYC, mTORC1 and unfolded-protein-response programmes, together with the heat-shock genes *HSPA1A*, *HSPA8* and *HSP90AA1* (Fig. 2g) ^9^. The predicted profiles captured a coordinated stress and proteostasis signature at both programme and gene levels. These signed means measure expression-effect fidelity rather than enrichment or pathway activation, and the selected case study does not establish a drug mechanism.

Aggregate gene and programme scores can remain accurate even when generated cells mis-represent the occupancy of response states. We therefore examined a selected Goserelin dose series in HCT15 cells to determine whether the generated population preserved response-state occupancy (Fig. 2c). The held-out 0.5 *µ*M condition, represented by 500 cells, lay between training doses of 0.05 and 5.0 *µ*M, each also represented by 500 cells. Its observed cell-cycle composition nevertheless lay outside the range spanned by the two endpoints: the G1 fraction was 49.6%, compared with 60.4% and 58.2% at the training doses, whereas the G2/M fraction was 37.6%, compared with 29.2% and 30.4%. Neither endpoint occupancy nor any mixture of the two reproduced the observed intermediate-dose population shift. In a shared UMAP, PerturbLDM more closely reproduced the occupancy of the observed cell-state regions than CPA or chemCPA. Both distributional distances showed the same ordering, although the numerical margins were modest: MMD-RBF was 0.2106, compared with 0.2396 for CPA and 0.2150 for chemCPA, and optimal-transport (OT) Wasserstein distance was 62.14, compared with 63.24 and 62.42, respectively. Absolute-expression metrics followed the same ordering (Supplementary Table S1). This comparison identified a population shift not captured by aggregate summaries, while remaining a selected illustration rather than benchmark-wide evidence.

We next assessed this distributional pattern across the full benchmark. For each held-out condition, we compared 500 observed and 500 generated cells using MMD-RBF and OT Wasserstein distance (Fig. 2h). MMD-RBF measures kernel-based discrepancy between two cell samples, whereas OT Wasserstein distance summarises the cost of matching one empirical distribution to the other. Lower values indicate closer local alignment. The two metrics produced a highly consistent ordering: across all 13,942 paired conditions, PerturbLDM had lower MMD-RBF and OT Wasserstein distances than both CPA and chemCPA in more than 99.9% of conditions in all four comparisons. Mean MMD-RBF was 0.2063 for PerturbLDM, 0.2426 for CPA and 0.2197 for chemCPA, whereas mean OT Wasserstein distance was 58.85, 60.26 and 59.43, respectively. The median paired MMD-RBF reductions were 0.03489 versus CPA (95% bootstrap CI, 0.03477–0.03499) and 0.01138 versus chemCPA (0.01128–0.01146). Corresponding OT Wasserstein reductions were 1.357 (1.353–1.361) and 0.5010 (0.4965–0.5059). BH-adjusted *P* was below 10*^−^*^300^ for all four two-sided paired Wilcoxon comparisons. The consistent reductions extended the comparison beyond aggregate response recovery to condition-aligned empirical distributions. MMD and OT assess local sample concordance but do not establish calibration of the full conditional response distribution.

The Tahoe-100M benchmark assessed three levels of response fidelity beyond absolute-profile reconstruction across 13,942 held-out conditions: matched-control-relative gene effects, coordinated signed programmes and condition-aligned local cell-state distributions. PerturbLDM improved each level relative to the corresponding comparators (Supplementary Tables S4 and S5).

### Tahoe-derived profiles nominate pharmacological relationships

Following quantitative evaluation in Tahoe-100M, we examined downstream use of precomputed response profiles generated by a Tahoe-trained frozen-SMILES variant in the PANACEA pharmacological assay^23^. The analysis evaluated whether Hallmark effects computed after control-only assay calibration recovered known pharmacological neighbourhoods in pathway space^1,2^.

Compounds with related pharmacology can converge on coordinated cellular programmes even when their individual gene-level responses differ. We therefore represented each of the 64 predicted PANACEA responses by its matched-control-relative Hallmark effects and ranked compounds by programme similarity. The resulting rankings were evaluated against independent Broad Drug Repurposing Hub MoA annotations ^26^ (Fig. 3a and Supplementary Fig. S6a). Among 27 evaluable drugs, 14 had a top-ranked candidate sharing at least one MoA annotation (51.9%, Wilson 95% CI, 34.0–69.3%), compared with 39.6% expected under the query-specific null (Fig. 3b and Supplementary Fig. S6d). The observed rate was therefore 12.3 percentage points above expectation (exact one-sided Poisson-binomial *P* = 0.1207) ^27,28^. Although this difference was not statistically significant, the same directional pattern was observed in both descriptive exact-identity strata: top-1 concordance reached 60.0% (6 of 10) and 47.1% (8 of 17) for drugs recorded and not recorded in the Tahoe training data, respectively, compared with corresponding expectations of 42.7% and 37.8% (Fig. 3b). Shared-MoA candidates also appeared early in both strata, with median first-positive ranks of 1 and 2, respectively (Fig. 3c and Supplementary Table S2).

**Fig. 3.**
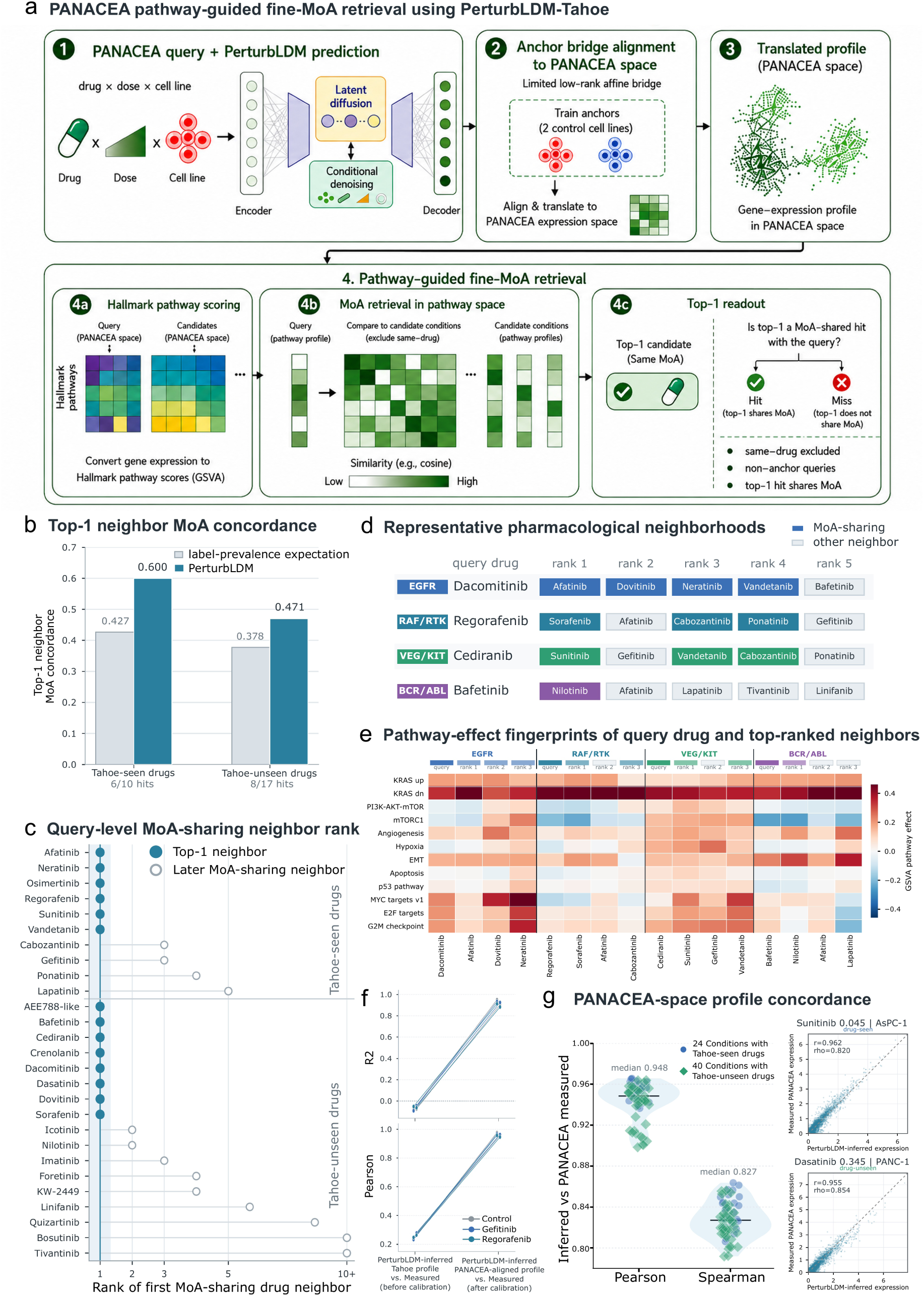
— Programme-level pharmacological neighbourhoods from Tahoe-derived response profiles. a, PANACEA application workflow. Precomputed profiles generated by a Tahoe-trained frozen-SMILES variant were calibrated with matched cell-line controls, converted to matched-control-relative Hallmark pathway effects and ranked by similarity after same-drug exclusion (Supplementary Fig. S6a and Supplementary Table S5). **b,** Descriptive top-1 MoA concordance stratified by exact drug-identity occurrence in the Tahoe training records (labelled Tahoe-seen and Tahoe-unseen in the panel). This was not a prospective drug-held-out split. Grey bars show query-specific label-prevalence expectations. The combined result was 14 of 27 (51.9%, Wilson 95% CI, 34.0–69.3%) and was not significant under the one-sided query-specific Poisson-binomial null (*P* = 0.1207). **c,** Rank of the first candidate sharing a Broad Hub MoA annotation with each of the 27 evaluable query drugs. **d,** Representative pharmacological neighbourhoods. Coloured boxes mark candidates sharing at least one MoA annotation with the query. **e,** Matched-control-relative Hallmark effects for the selected queries and their three highest-ranked neighbours, showing coordinated programme-level similarities underlying the rankings. **f,** Expression-profile concordance for six shared profiles (two controls and four treatments) before and after control-only calibration. Lines connect matched profiles. Calibration summaries are descriptive. **g,** PANACEA-space profile concordance across 64 grouped conditions (24 with and 40 without exact drug identity in the Tahoe records). The main display shows Pearson and Spearman summaries. Insets show two illustrative condition-level expression scatterplots. Across the same conditions, median *R*^2^ was 0.893.

To identify the pharmacological relationships underlying the aggregate ranking pattern, we examined four representative top-ranked neighbourhoods. Dacomitinib ranked Afatinib first, consistent with their EGFR/ERBB pharmacology^29,30^, whereas Regorafenib ranked Sorafenib within an overlapping RAF, KIT and VEGFR context ^31,32^. Cediranib ranked Sunitinib within VEGFR-directed pharmacology^33,34^. Bafetinib similarly ranked Nilotinib within the BCR–ABL inhibitor class ^35,36^ (Fig. 3d). Correlations between their matched-control-relative Hallmark vectors ranged from 0.9746 to 0.9867, indicating aligned directions of change across multiple response programmes (Fig. 3e). Across annotations, top-1 concordance was highest for EGFR, KIT, PDGFR and RAF terms and lower for rarer or more pharmacologically heterogeneous terms (Supplementary Fig. S6f). Because these examples were selected after ranking, they show how programme similarity aligned with known pharmacology but do not provide independent validation or MoA discovery.

The predicted and measured profiles originated from different assay settings, motivating a separate evaluation of alignment. Across six shared profiles, comprising two controls and four treatments, control-only calibration increased median *R*^2^ from -0.069 to 0.907 and median Pearson correlation from 0.256 to 0.955 (Fig. 3f). For the full set of 64 grouped PANACEA conditions, the calibrated profiles reached median Pearson *r* = 0.948 and Spearman *ρ* = 0.827 in the displayed summaries. The corresponding median *R*^2^ was 0.893 (Fig. 3g). These concordance values supported pathway-neighbourhood analysis of the calibrated profiles but did not independently validate the generative model. The combined ranking and assay-alignment results illustrate how Tahoe-derived response profiles can be repurposed for pharmacological neighbourhood analysis (Supplementary Table S5).

### Recovering a mid-gestational fetal-colon state

Intermediate intestinal development involves coordinated changes in progenitor, absorptive and secretory epithelial programmes rather than uniform interpolation between observed endpoints ^24^. We therefore trained PerturbLDM on early (PCW 11–14, broad stage 1) and mature (PCW 19–23, broad stage 3) enterocytes and generated the withheld mid-gestational population of 1,529 PCW 16–18 cells at broad stage 2 (Fig. 4a). This endpoint-to-intermediate design evaluated whether conditional generation could recover the transcriptional organisation of the intervening population rather than default to either observed anchor.

**Fig. 4.**
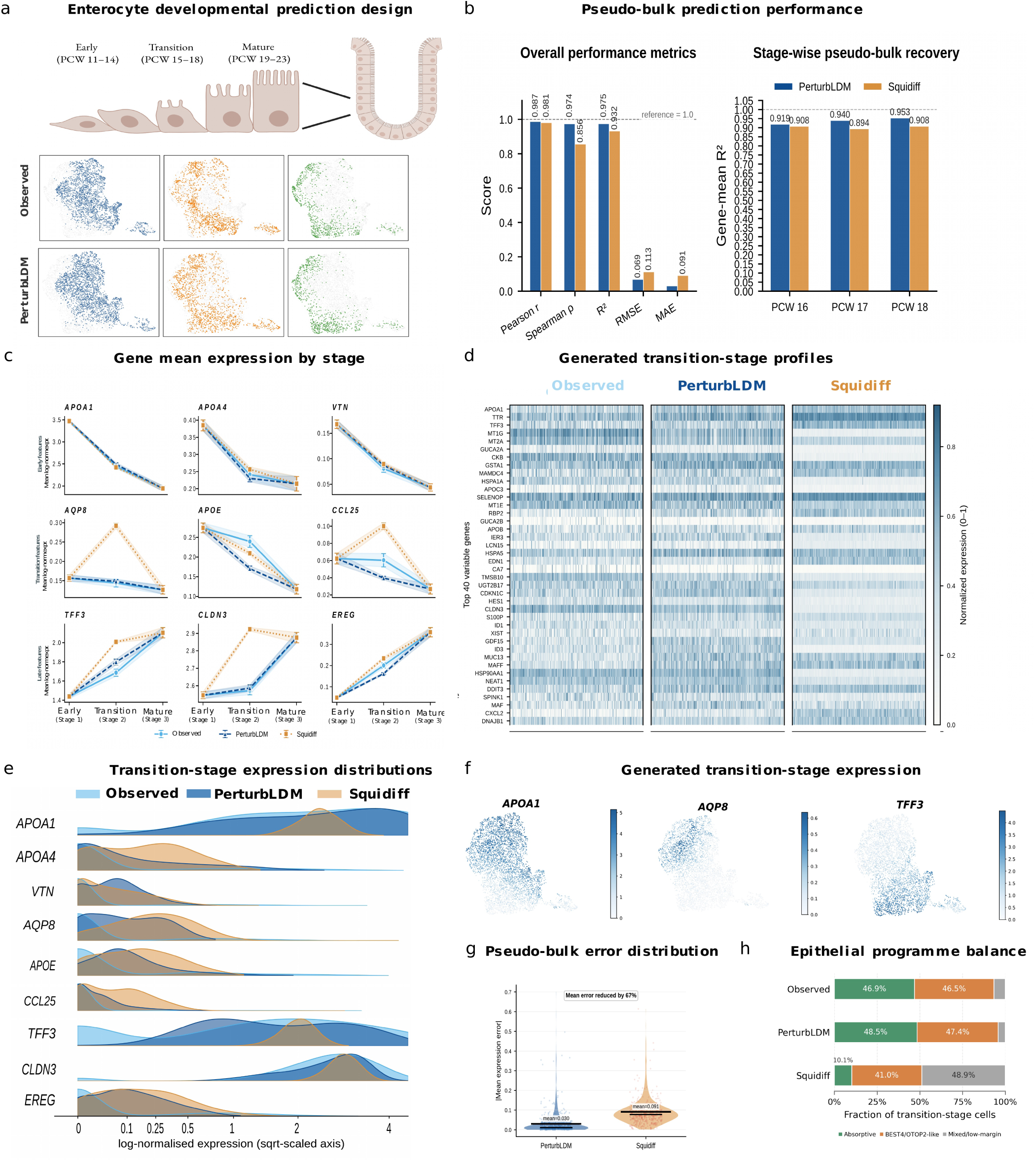
— Conditional generation of an intermediate developmental state in human fetal colon. a, Experimental design and shared reference UMAPs arranged by broad developmental stage. Columns show early, transition and mature populations. The upper and lower UMAP rows show observed and PerturbLDM-generated cells, respectively. Early PCW 11–14 and mature PCW 19–23 enterocytes served as observed anchors. The transition population was generated using the pooled broad-stage-2 condition and evaluated against 1,529 held-out PCW 16–18 cells. **b,** Pooled pseudo-bulk Pearson correlation, Spearman correlation, *R*^2^, RMSE and MAE across the shared 800-HVG feature space. The model treated PCW 16–18 as one pooled broad-stage-2 transition condition. The adjacent gene-mean *R*^2^ values show a descriptive stratification by annotated PCW. **c,** Observed and predicted stage-wise mean expression (*±*s.e.m. across cells) for nine epithelial genes selected retrospectively using published annotations and prespecified recovery criteria across early (stage 1), transition (stage 2) and mature (stage 3) populations: *APOA1*, *APOA4*, *VTN*, *AQP8*, *APOE*, *CCL25*, *TFF3*, *CLDN3* and *EREG*. The transition population was withheld for evaluation. Curves compare observed profiles with PerturbLDM and Squidiff predictions. Error bars are descriptive within-population summaries of nested cells and do not represent donor-level uncertainty. **d,** Normalised-expression heatmaps for the 40 most variable genes across 1,529 observed, PerturbLDM-generated and Squidiff-generated transition-stage cells, comparing within-population expression variation. Cells were not paired across groups. **e,** Kernel-density distributions of log-normalised single-cell expression for the same nine genes as in **c** across the held-out PCW 16–18 population, including zero-valued cells. The display axis is square-root-scaled. **f,** Expression maps of *APOA1*, *AQP8* and *TFF3* across PerturbLDM-generated transition-stage cells. Each gene uses its own expression scale. **g,** Distribution of per-gene absolute differences between predicted and observed pseudo-bulk means across 800 HVGs on the log-normalised expression scale. Mean errors were 0.030 for PerturbLDM and 0.091 for Squidiff, corresponding to a 67% reduction. Additional feature-level and PCW-stratified summaries are reported in Supplementary Table S6. **h,** Exploratory proportions of observed, PerturbLDM-generated and Squidiff-generated transition-stage cells assigned to absorptive-trajectory, BEST4/OTOP2-like or mixed/low-margin epithelial programmes. Gene-wise standardisation parameters were estimated from observed cells across all three broad stages, and the resulting parameters and score-margin rule were applied unchanged to all profiles. Percentages are descriptive cell-level summaries.

To match the feature-space scale of published Squidiff examples, we evaluated both models on the same set of 800 highly variable genes (HVGs) ^21^. Both methods captured the pooled transition-state profile, but PerturbLDM more closely preserved gene ordering and expression magnitude. Against the pooled PCW 16–18 target, PerturbLDM achieved Pearson *r* = 0.9873, Spearman *ρ* = 0.9739 and *R*^2^ = 0.9746, compared with 0.9806, 0.8561 and 0.9317 for Squidiff, respectively. Root-mean-square error was 0.069 versus 0.113, and MAE was 0.030 versus 0.091 (Fig. 4b,g), corresponding to a 67% reduction in mean per-gene error.

To examine within-population variation, we placed observed and generated cells in a shared reference embedding. PerturbLDM-generated cells occupied regions similar to the observed PCW 16–18 population (Fig. 4a). Although PCW 16–18 cells were generated under one pooled transition condition, descriptive stratification by annotated PCW showed consistently higher gene-mean *R*^2^ for PerturbLDM than for Squidiff at PCW 16, 17 and 18, with values of 0.919–0.953 and 0.894–0.908, respectively (Fig. 4b). Variable-gene heatmaps and single-gene density estimates showed cell-to-cell expression variation across the generated transition population (Fig. 4d,e). An exploratory analysis compared a broad absorptive maturation trajectory with the transcriptionally distinct BEST4/OTOP2 programme established early in intestinal development ^24^. Absorptive-trajectory and BEST4/OTOP2-like assignments comprised 48.5% and 47.4% of PerturbLDM-generated cells, compared with 46.9% and 46.5% of observed cells. Squidiff assigned 10.1% of cells to the absorptive programme and 48.9% to the mixed or low-margin group, consistent with weaker separation of the two programme scores (Fig. 4h).

Developmental interpretation focused on epithelial programmes with distinct behaviour across gestation. Nine stage-resolved genes represented absorptive and lipid-handling features (*APOA1* and *APOA4*), the early progenitor-associated feature *VTN*, the maturation-associated transport feature *AQP8*, regional epithelial features (*APOE* and *CCL25*) and secretory, barrier or epithelial-signalling features (*TFF3*, *CLDN3* and *EREG*)^24^ (Fig. 4c,f). These genes spanned decreasing, increasing and non-monotonic stage-associated trajectories, allowing the predicted transition state to be assessed against coordinated rather than uniform expression changes. Across all 800 HVGs, the median per-gene absolute difference between predicted and observed pseudo-bulk means was 0.0107 for PerturbLDM and 0.0774 for Squidiff, and 745 of 800 genes (93.1%) versus 547 (68.4%) fell within a descriptive absolute-error tolerance of 0.1 on the log-normalised scale (Supplementary Fig. S7 and Supplementary Table S6). The pooled profile, stage-associated epithelial features, cell-level displays and gene-wide errors collectively supported closer recovery of a mid-gestational enterocyte-like population than Squidiff.

### Predicting lineage-specific IFN-*β* responses

Type I interferon induces a shared antiviral programme, but its magnitude and accompanying inflammatory programmes vary across lineages and cellular states^37,38^. We evaluated whether IFN-*β* response programmes learned across profiled myeloid and lymphoid lineages generalised to withheld stimulated states within each compartment. In the Kang et al. dataset^25^, training included both control and stimulated CD4 T cells, NK cells, CD14^+^ monocytes and dendritic cells. Stimulated B cells, CD8 T cells and FCGR3A^+^ monocytes were withheld while their control cells were retained (Fig. 5a). We compared PerturbLDM with scGen^15^ using stimulated-state whole-profile concordance, matched-control DEGs and GO programmes.

**Fig. 5.**
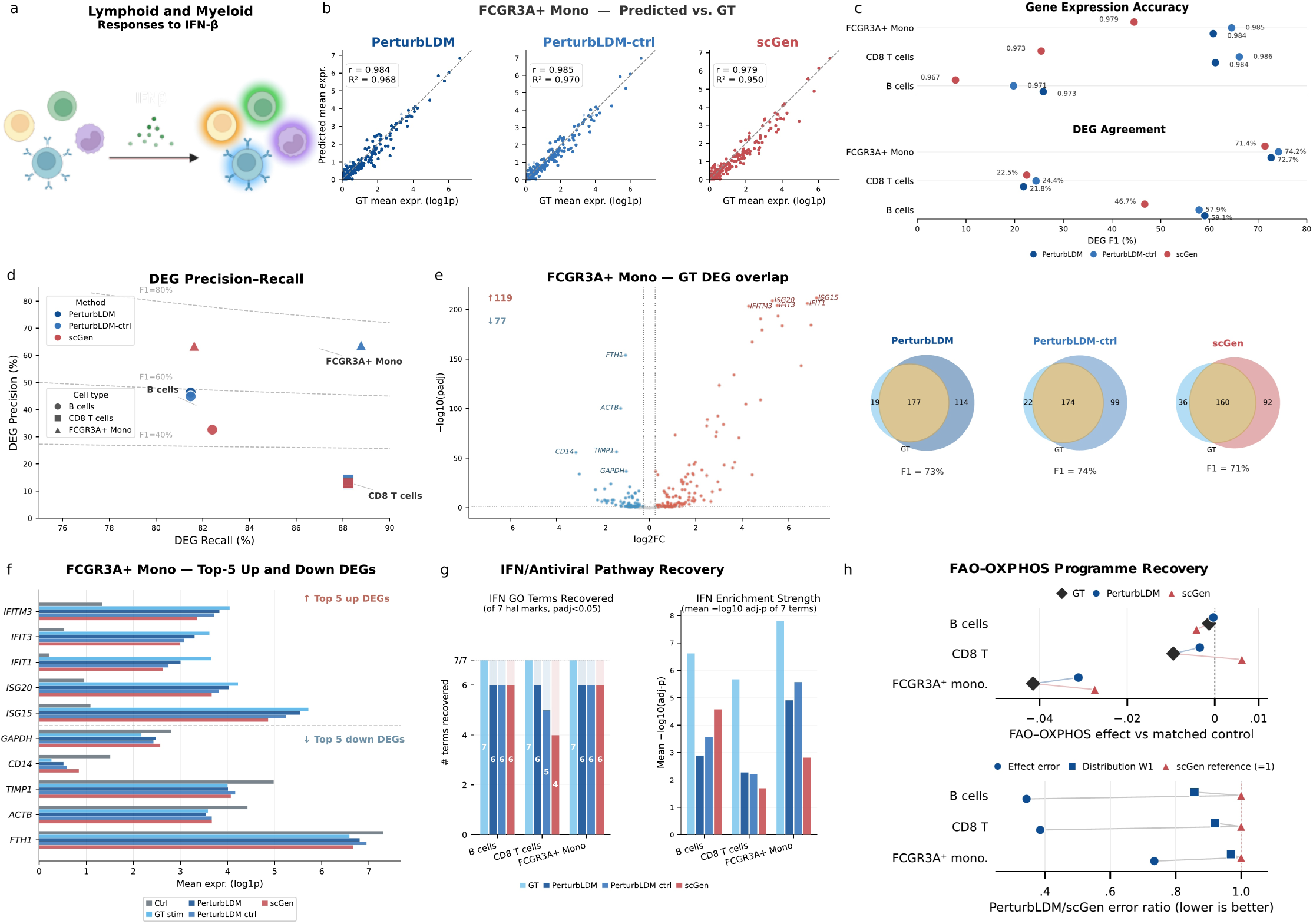
Conditional generation of lineage-specific IFN-*β* responses in PBMCs. **a,** Lineage-specific IFN-*β* stimulation. Stimulated B cells, CD8 T cells and FCGR3A^+^ monocytes were withheld, while their lineage-matched unstimulated controls were retained. **b,** Selected FCGR3A^+^-monocyte illustration of gene-mean expression concordance for PerturbLDM, PerturbLDM-ctrl and scGen. PerturbLDM-ctrl denotes a control-initialised sensitivity analysis, with the displayed strength selected against the held-out targets. **c,** Lineage-level stimulated-state gene-mean Pearson correlation and matched-control DEG F1 for the three methods. Cross-lineage means are unweighted descriptive summaries (*n* = 3 lineages). **d,** Lineage-level DEG precision–recall relationships with iso-F1 guides. **e,** Ground-truth FCGR3A^+^-monocyte DEG volcano plot and method-versus-ground-truth overlap diagrams based on bidirectional DEG calls. Ground-truth calls compare measured stimulated cells with matched measured controls. Prediction calls compare each predicted stimulated profile with the same matched controls. DEGs were called at adjusted *P <* 0.05 and *|* log_2_ FC*| >* 0.25. These within-profile tests do not provide model-comparison inference. The analysis in **g** used separately defined upregulated-gene lists rather than these bidirectional overlap-call sets. **f,** Mean expression of the five most upregulated (*IFITM3*, *IFIT3*, *IFIT1*, *ISG20* and *ISG15*) and five most downregulated (*GAPDH*, *CD14*, *TIMP1*, *ACTB* and *FTH1*) ground-truth FCGR3A^+^-monocyte DEGs across control, ground truth and predictions. **g,** Recovery count and mean *−* log_10_ adjusted enrichment *P* value for seven GO Biological Process terms selected from ground-truth enrichment results spanning interferon responses, antiviral defence and regulation of viral processes. Enrichment used separately recomputed upregulated-gene lists rather than the bidirectional overlap calls in **e**. These are selected within-profile descriptive summaries rather than method-level tests. **h,** Lineage-resolved recovery of an FAO–OXPHOS transcriptional programme. Top, matched-control-relative composite effects for measured stimulated profiles, PerturbLDM and scGen across the three held-out lineages. Points are vertically offset within each lineage for visibility.The composite is the arithmetic mean of independently computed FAO and OXPHOS scores. Bottom, PerturbLDM error divided by scGen error for the composite effect (blue circles) and the per-cell composite-score distribution (blue squares, one-dimensional Wasserstein distance). Red triangles mark the scGen reference at 1. Values below 1 favour PerturbLDM. All six ratios were below 1. The FAO and OXPHOS scores used 7 and 31 genes, respectively, represented in the shared 2,000-HVG space. Scores reflect programme-associated gene expression rather than metabolic flux.

PerturbLDM and its control-initialised inference variant, PerturbLDM-ctrl (Methods), more accurately reproduced the complete stimulated-state profiles of the three held-out cell types than scGen. The two variants produced nearly identical mean Pearson correlations (0.9804 and 0.9807), Spearman correlations (0.9084 and 0.9087), *R*^2^ values (0.9593 and 0.9603) and MAEs (0.0367 and 0.0363), compared with 0.9731, 0.8371, 0.9429 and 0.0486 for scGen, respectively (Fig. 5c, Supplementary Fig. S9 and Supplementary Table S6). In the displayed FCGR3A^+^-monocyte example, PerturbLDM-ctrl reached Pearson *r* = 0.9852 and *R*^2^ = 0.9699, compared with 0.9789 and 0.9502 for scGen (Fig. 5b). The similar predictions obtained from Gaussian and control-based initialisation were consistent with the conditional denoiser guiding both tested starting states towards similar lineage-specific stimulated profiles. Because the control-initialisation strength was selected against the same held-out targets, this comparison was treated as a sensitivity analysis.

Matched-control DEG analysis assessed intervention-specific recovery by comparing predicted and observed IFN-*β*-responsive DEGs relative to matched controls. Across the three held-out cell types, mean DEG F1 was 51.2% for PerturbLDM and 52.2% for PerturbLDM-ctrl, compared with 46.9% for scGen. Corresponding mean precisions were 39.9%, 40.9% and 36.3%, respectively (Fig. 5c–e). PerturbLDM showed the largest improvement in B cells, with an F1 of 59.1% versus 46.7% for scGen. DEG F1 values were similar in CD8 T cells for PerturbLDM and scGen (21.8% and 22.5%), whereas PerturbLDM-ctrl reached 24.4%. In FCGR3A^+^ monocytes, PerturbLDM-ctrl reached an F1 of 74.2% versus 71.4% for scGen with a smaller predicted DEG set (492 versus 748 genes). The observed response was led by canonical interferon-stimulated genes, including *IFITM3*, *IFIT3*, *IFIT1*, *ISG20* and *ISG15* (Fig. 5f).

To test whether gene-level recovery preserved coordinated antiviral biology, we examined seven ground-truth-derived GO Biological Process terms spanning interferon responses, antiviral defence and regulation of viral processes. PerturbLDM recovered 6 of 7 terms in each held-out cell type. scGen recovered 6 of 7 terms in B cells and FCGR3A^+^ monocytes but 4 of 7 in CD8 T cells, whereas PerturbLDM-ctrl recovered 6 of 7 in B cells and monocytes and 5 of 7 in CD8 T cells (Fig. 5g). The broader monocyte comparison yielded a top-20 GO-term Jaccard similarity of 0.667 for PerturbLDM-ctrl and 0.290 for scGen (Supplementary Fig. S10). Acute type I interferon signalling can promote a coupled fatty-acid oxidation (FAO)–oxidative phosphorylation (OXPHOS) programme, with FAO providing substrates for mitochondrial respiration ^39,40^. We therefore examined whether the predicted profiles captured this lineage-resolved transcriptional programme, summarised as the equal-weight mean of independently computed FAO and OXPHOS scores. Across the three held-out lineages, PerturbLDM predictions deviated 27–66% less from the measured FAO–OXPHOS composite effects than scGen predictions (Fig. 5h and Supplementary Fig. S13). Per-cell programme-score distributions showed the same pattern, with Wasserstein distances to the measured distributions reduced by 3–14% relative to scGen. Together, these analyses showed recovery of canonical antiviral programmes and an interferon-associated FAO–OXPHOS transcriptional programme.

## Discussion

PerturbLDM applies conditional latent diffusion to generate single-cell transcriptional responses for unmeasured conditions across biological contexts and data scales. We evaluated absolute-expression fidelity alongside intervention-induced gene effects, coordinated programmes and local population structure. Tahoe-100M supplied the condition coverage for model fitting and the central benchmark, and PANACEA evaluated downstream reuse of Tahoe-derived response profiles. Separately fitted fetal-colon and PBMC models tested the framework in developmental and immune settings.

PerturbLDM represents biological conditions with different structures by integrating categorical identities, continuous covariates and expression-defined cellular contexts within a common conditional-denoising framework. In Tahoe, these forms correspond to a learned drug embedding, a log_10_-scaled dose scalar that preserves order-of-magnitude spacing and the matched-control expression profile. This modular design could accommodate further conditions, such as treatment duration or disease state, as sufficiently annotated datasets become available. The Tahoe random split held out new combinations at the three observed dose levels, thereby evaluating recombination rather than interpolation to unmeasured doses. The conditioning pathway can accept other positive dose values and transform them nonlinearly, but generation was evaluated only at 0.05, 0.5 and 5.0 *µ*M. In fetal colon, stages 1, 2 and 3 define an ordered scalar coordinate. The stage value is transformed by an MLP before conditioning reverse diffusion, making the generated stage 2 profile a nonlinear function of this coordinate rather than an average of the endpoint expression or latent representations. By contrast, Squidiff constructs the intermediate semantic condition by linearly interpolating the endpoint latents, equivalent at stage 2 to their midpoint average ^21^. These examples distinguish recombination of observed conditions from interpolation along an ordered coordinate and show how condition construction determines the generalisation being tested.

Recent benchmarks have shown that simple baselines can remain competitive with learned models in perturbation-response prediction ^12,13,41^. We observed a similar pattern in Tahoe, where AdditiveMean captured much of the absolute-expression variation (Fig. 2b). Because stable cellular background can inflate absolute-expression metrics, AdditiveMean’s combination of cell-line and drug-dose marginals provided a direct test of whether PerturbLDM captured their context-specific interaction. Matched-control effect Pearson correlation was higher for PerturbLDM in 95.2% of the 13,942 held-out conditions, and PerturbLDM also reproduced Hallmark programme effects more accurately than AdditiveMean, indicating that the advantage extended from background-preserving profiles to intervention-induced gene and programme responses. For more than 99.9% of conditions, PerturbLDM also yielded lower condition-aligned MMD and OT discrepancies than CPA and chemCPA, indicating closer empirical cell-state distributions within matched conditions. The Goserelin dose series illustrated this distributional distinction through a dose-specific shift in cell-cycle occupancy, whereas the selected Bortezomib condition showed concordant predicted and observed effects across stress, proteostasis and heat-shock responses. Tahoe therefore provides evidence that conditional latent diffusion can recover response structure beyond separable factor averages, within the defined space of drugs, doses and cell lines observed during training.

To examine whether Tahoe-derived response profiles remained informative across a different experimental assay, we applied them to PANACEA ^9,10,23,42,43^. At the gene-response level, control-only assay calibration aligned Tahoe-derived predictions with the PANACEA expression space, enabling comparison of condition-specific transcriptional responses. At the drug-action level, converting those profiles to matched-control-relative Hallmark effects produced pharmacological neighbourhoods containing several established inhibitor relationships. Top-1 MoA concordance exceeded the query-specific expectation in both exact-identity strata, although the combined result did not reach significance under the query-specific null. At both levels, assay-aligned atlas predictions nominated drug-context conditions and pharmacological relationships for targeted experimental follow-up.

Comparably scaled perturbation data are not available for every biological system. We therefore fitted PerturbLDM directly to the smaller fetal-colon and PBMC datasets, tailoring each analysis to the developmental or immune-response question. In fetal colon, PerturbLDM more closely matched the withheld mid-gestational profile than Squidiff and preserved stage-associated changes across progenitor, absorptive, regional and secretory epithelial features. It also retained the observed balance between absorptive and BEST4/OTOP2-like epithelial programmes more faithfully than Squidiff, extending the comparison to within-population developmental organisation^24^. The generated transition state also reproduced increasing, decreasing and non-monotonic trajectories across representative epithelial genes (Fig. 4c), extending the comparison beyond the pooled profile. In PBMCs, responses learned across lymphoid and myeloid cell types supported generation of withheld IFN-*β*-stimulated states: the complete stimulated profiles were reproduced more accurately than by scGen, interferon and antiviral programmes were captured across lineages, and DEG-level fidelity varied by lineage. PerturbLDM also reproduced lineage-resolved effects in the interferon-associated FAO–OXPHOS transcriptional programme with lower error than scGen, extending the biological comparison beyond canonical interferon-stimulated genes. In plasmacytoid dendritic cells, type I interferon signalling increases FAO and OXPHOS partly through PPAR*α*. This metabolic shift supports the bioenergetic demands of activation and contributes to antiviral protection ^39^. At the transcriptomic level, the reproduced FAO–OXPHOS programme indicates that the generated profiles retained coordinated metabolic gene expression associated with the stimulated state. Together, these applications showed that more accurate generated states retained interpretable developmental, antiviral and metabolic programmes.

Perturbation-response prediction can be viewed as a biological state-forecasting problem, conceptually related to next-state prediction in learned world models and the broader virtual-cell programme ^14,43^. Rather than a deterministic next frame, the target is a conditional distribution of molecular states after intervention, encompassing coordinated gene programmes, state occupancy, multimodality and cell-to-cell variability. Progress towards this objective will depend on biological coverage as well as cell count, including additional perturbation classes and combinations, dose and time courses, tissues, donors and paired temporal or multimodal measurements. How coverage and model capacity jointly shape generalisation remains to be tested across data compositions ^44^. Relevant evaluations include marginal baselines, factor-heldout and cross-dataset tests, prospective experiments and uncertainty calibration. The MMD and OT analyses in this study quantify local distributional concordance without establishing global calibration of the predicted conditional distributions. The Tahoe analyses indicate that explicit distribution-aware objectives may improve the modelling of cellular population structure beyond mean responses. Such objectives could be paired with metrics that assess the calibration of population composition and within-state variability. Biological priors may also help extend conditional generation to genetic and combinatorial interventions. Together, these extensions could support probabilistic forecasts of cellular outcomes, but predictive fidelity would remain distinct from causal identification of gene-regulatory mechanisms^17,41^.

## Methods

### Model overview

PerturbLDM generated condition-specific responses by combining a variational autoencoder with conditional diffusion. The autoencoder mapped normalised expression profiles **x** ∈ R*^G^* to compact latent cell states **z**_0_ ∈ R*^Dz^* ^45^. A denoising model then sampled latent states under biological condition *c*, and the decoder mapped these states back to gene expression. We use the term transcriptomic manifold to denote this learned latent space rather than a directly measured biological axis.

The autoencoder defined the representation space. The diffusion model generated context-conditioned latent variation, and the conditioning interface specified the requested response. Depending on the task, *c* encoded drug identity, dose, matched-control expression, developmental stage, stimulation state or other categorical labels. Modelling diffusion in the latent space reduced the dimensionality of the generative problem^22,46^. The complete architecture is shown in Supplementary Fig. S1.

### Latent transcriptomic autoencoder

We learned the latent representation with a multilayer-perceptron (MLP) variational autoencoder ^45^. The encoder mapped **x** to the parameters of a diagonal Gaussian posterior,

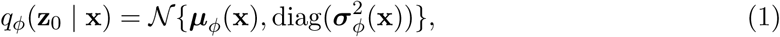

and sampled latent states with the reparameterisation trick during training. The decoder *d_θ_*mapped **z**_0_ back to expression space. Encoder and decoder modules were multilayer perceptrons with batch normalisation, ReLU activation and dropout. Unless otherwise stated, the decoder used a Gaussian reconstruction model with mean-squared reconstruction loss.

The autoencoder objective was

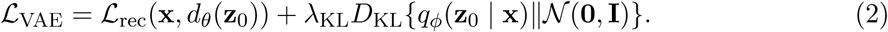

After autoencoder training, we encoded training cells without sampling noise and retained the posterior mean latent code. These cached latents served as clean targets for diffusion training.

### Latent diffusion model

The latent diffusion component generated responses by recovering clean latent cell states from progressively noised samples under condition *c* ^46^. We indexed latent states as **z**_0_*, …,* **z***_T_*, where **z**_0_ is clean and **z***_T_* is terminal noise. The equations use *t*= 1*, …, T*, with *T* = 1000 diffusion transitions, whereas the implementation stored the corresponding scheduler indices as 0*, …, T* −1. The forward process added Gaussian noise as

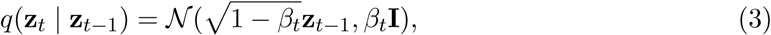

or equivalently

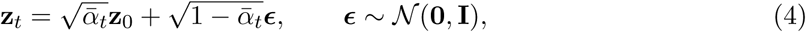

where 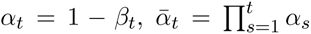 and *ᾱ*_0_ = 1. A DDPM scheduler with a linear variance schedule from *β*_1_ = 0.00085 to *β_T_* = 0.015 was used. The scheduler clipping range was set to the maximum absolute value of the cached training latents for each dataset. This clipping was used only to stabilise scheduler updates in latent space and did not modify cached training latents, decoded ground-truth profiles or evaluation targets.

The learned reverse process is a conditional Gaussian transition,

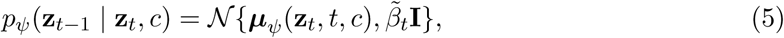

With

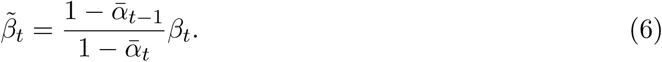

The denoising network *g_ψ_* receives the noisy latent **z***_t_*, the corresponding zero-based scheduler index *s* = *t* − 1 and condition *c*, and outputs **ŷ***_t_* = *g_ψ_*(**z***_t_, s, c*). The implementation supports three standard prediction parameterisations ^46,47^, with training target

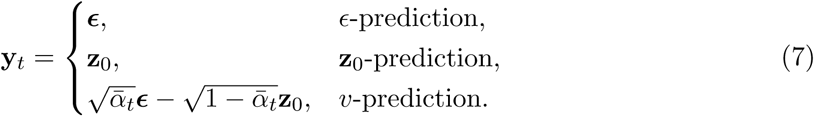

The denoiser was trained by mean-squared error against the selected target,

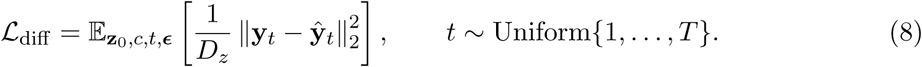

The scheduler converts the selected target parameterisation into a clean-latent estimate,

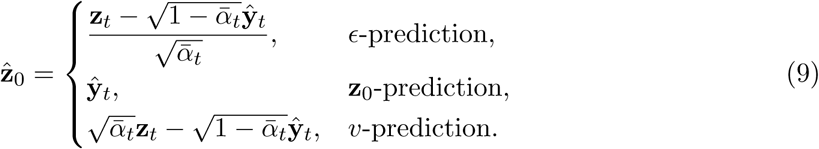

After clipping z^0 to the configured latent range, the DDPM posterior mean is

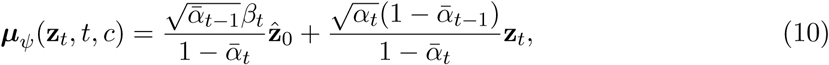

and one reverse step is sampled as

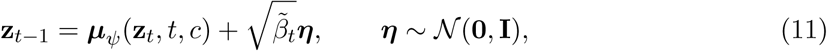

with the stochastic term omitted at the final step. Tahoe, colon development and PBMC used *v*-prediction.

### Conditional denoising networks

Conditioning was applied during denoising so that biological context and diffusion time jointly determined each reverse step. PerturbLDM used residual MLP denoisers with a shared latent backbone and task-specific conditioning interfaces. Biological conditions and diffusion time remained separate inputs until their fusion in the residual backbone.

**Condition encoder.** For mixed categorical and continuous conditions, each of the *M* condition fields was first mapped to a shared context dimension. For *j* = 1*, …, M*,

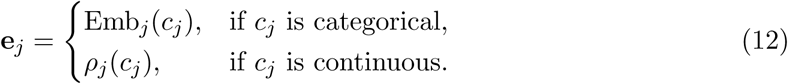

where *ρ_j_* is a linear projection for vector-valued covariates and a small SiLU MLP for scalar covariates. The condition descriptor is

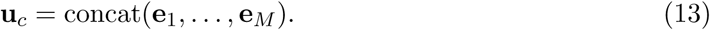

**Timestep encoder.** Diffusion transition indices were represented by sinusoidal embeddings. Let *s* = *t* − 1 be the zero-based scheduler index and *m* = [*d_t_/*2♩, where *d_t_* is the timestep-embedding dimension. For *k* = 0*, …, m* − 1,

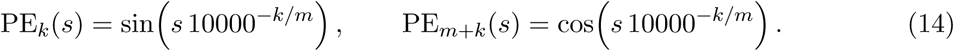

The embedding was zero-padded when *d_t_* was odd and projected into the context space,

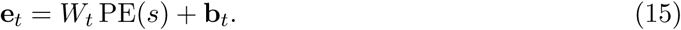

**Condition-time fusion and residual denoising.** The condition descriptor and timestep embedding are concatenated and mapped to the denoiser hidden dimension,

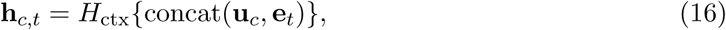

where *H*_ctx_ is a two-layer MLP with batch normalisation and SiLU activation.

The noisy latent is projected as

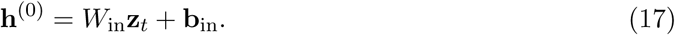

At residual block *l*, the condition embedding is added to the hidden state before the block MLP,

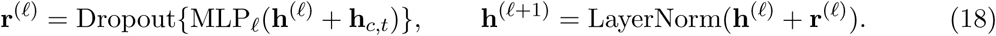

Each MLP*_l_*comprised two linear layers separated by a SiLU activation. The final block output was mapped back to latent dimensionality to produce **ŷ***_t_*.

The Tahoe drug-response model used a dedicated drug-dose denoiser. Drug identity was represented by a learned embedding, dose was represented as a scalar dose feature, and matched DMSO control expression from the same cell line was projected into the context space. These context terms and the timestep embedding were fused by the same context-encoder and residual-MLP design. Colon development and PBMC used the generic mixed-condition denoiser. Supplementary Fig. S2 reports single-run sensitivity analyses of Tahoe implementation variants. These analyses were not designed as replicated ablations.

We use *Tahoe pretraining* to denote fitting the Tahoe drug-response model before its core held-out benchmark and intended downstream use. The PANACEA application uses precomputed profiles generated by a Tahoe-trained frozen-SMILES variant. Fetal colon and PBMC use task-specific fits of PerturbLDM, testing framework-level applicability rather than reuse of Tahoe-trained parameters.

### Training and inference

Training comprised two stages. We first trained the autoencoder on normalised expression and then trained the denoiser on cached latent codes. Both stages used AdamW, linear learning-rate decay and gradient clipping. At each denoiser update, we sampled a timestep uniformly, added Gaussian noise with the DDPM scheduler and optimised the model against the corresponding target.

Inference began with Gaussian latent noise and followed the learned reverse process for 1000 DDPM steps. We decoded the final latent sample and set negative expression values to zero before aggregation and metric calculation. Tahoe timestep-specific samples were retained for reverse-diffusion visualisation (Supplementary Fig. S4). Selected reverse-diffusion snapshots for fetal colon and PBMC are shown in Supplementary Figs. S8, S11 and S12. For PBMCs, we also evaluated a control-initialised variant, PerturbLDM-ctrl. We encoded control cells, added noise to an internal timestep and denoised under the target stimulation condition. Strengths from 0 to 1 were scored against the held-out stimulated targets. Strength 0.4 was selected retrospectively from this same-target sweep, and the resulting comparison was treated as a sensitivity analysis.

### Tahoe drug-response atlas

Tahoe supplied the data for model fitting and the primary conditional-generation benchmark. We normalised cells to 10,000 counts per cell and log-transformed the data. Genes detected in fewer than three cells were removed, and those detected in at least 1% of cells were retained. The resulting feature space contained 13,784 genes. Each evaluation unit comprised one drug, dose and cell line. Measured treatment conditions were withheld for direct comparison between predicted and observed responses (Supplementary Fig. S3a–d).

The Tahoe conditioning variables were drug identity, dose and matched control expression. The three dose levels (0.05, 0.5 and 5.0 *µ*M) were encoded as log_10_(*d/*0.05), where *d* is the dose in micromolar. Matched control expression was the mean normalised expression profile of DMSO-treated cells from the same cell line. Conditions with fewer than 500 cells were excluded, and larger conditions were capped at 500 cells for training and inference. During inference, each test condition was generated with 500 latent samples, decoded and averaged to obtain the condition-level prediction.

The Tahoe model used a 1,024-dimensional latent space, five residual blocks, hidden dimension 2,048, context dimension 768, timestep embedding dimension 256, dropout 0.1, batch size 512, learning rate 1 × 10*^−^*^4^ and four diffusion-training epochs. Training and inference used *v*-prediction.

### Tahoe baselines and effect analyses

We compared Tahoe predictions with learned models and training-set controls. Learned baselines included an MLP, a random forest, CPA and chemCPA^10,16^. Supplementary Methods provide implementation details. For these learned comparators, 3,252 conditions were reserved from the 32,529-condition development set for validation, leaving 29,277 fitting conditions. The same condition-level partition was used across methods, and the 13,942 held-out response conditions were used only for final evaluation. The controls quantified how much of each held-out response was explained by matched controls or train-set marginals. MatchedCtrl used the matched DMSO control profile as the prediction. CellLineMean predicted the train-set mean profile for the target cell line. DrugDoseMean predicted the train-set mean profile for the target drug-dose pair across cell lines. AdditiveMean combined train-only marginal means as

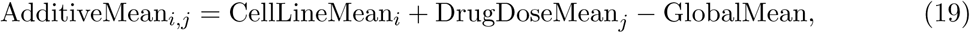

where *i*indexes the cell line and *j* indexes the drug-dose pair. AdditiveMean captures separable cell-line background and global drug-dose response^12,13^. Each marginal was an unweighted mean over training-condition profiles, and all marginal quantities were computed from the training split only.

For perturbation-effect analyses, observed and predicted effects were defined as 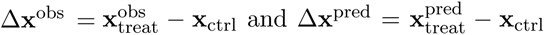, using the same matched DMSO control profile from the target cell line. Effect recovery was evaluated by all-gene effect Pearson correlation, all-gene effect Spearman correlation and effect MAE. Context-specificity analyses grouped conditions within fixed drug-dose strata or fixed cell-line-dose strata, and compared paired response-rank Spearman gain and centred-error reduction relative to AdditiveMean. MatchedCtrl was not used for effect-correlation baselines when the predicted effect vector was constant zero.

Signed Hallmark effect scores were computed as the mean matched-control-relative expression effect over member genes of each MSigDB Hallmark v2024.1 gene set^48^. These scores were used as signed gene-programme effect summaries, not as formal enrichment, GSVA or ssGSEA scores. Condition-level Hallmark profile concordance was evaluated by correlating predicted and observed signed Hallmark score vectors. Per-programme concordance was evaluated by correlating predicted and observed scores across held-out conditions for each Hallmark programme. Distributional analyses used condition-aligned predicted and observed single-cell samples.

The primary metrics were MMD-RBF^49^ and optimal-transport Wasserstein distance^50^. Energy distance and sliced Wasserstein distance were secondary analyses. Stratified Tahoe absolute-profile, effect- and Hallmark-level concordance summaries described variation across cell-line, drug and dose strata (Supplementary Fig. S5).

### Task-specific external comparators

scGen and Squidiff were task-specific comparators and were not included in the Tahoe benchmark. scGen was used for PBMC IFN-*β* stimulation transfer ^15^. Stimulated cells from selected lineages were withheld, and scGen inferred their profiles from learned control-to-stimulated shifts. Supplementary Methods describe the split, latent-shift construction and evaluation. Squidiff was used for fetal-colon maturation, where the target state lay between early and mature developmental anchors^21^. These task-specific comparisons were not pooled with the Tahoe benchmark.

### PANACEA response-programme neighbourhood analysis

#### PANACEA setting

The PANACEA analysis used a precomputed prediction matrix with 128 rows and 13,784 genes generated by a Tahoe-trained frozen-SMILES embedding variant. Condition metadata paired two rows per condition in their original order to produce 64 drug-dose-cell-line profiles. Downstream calibration, pathway scoring and ranking were performed on these profiles. For descriptive reporting, drugs were stratified by whether the exact identity occurred in the Tahoe training records. This label did not affect neighbour ranking. Analyses used 12,380 genes shared between the prediction matrix and PANACEA after gene-name harmonisation.

#### PANACEA-space calibration

Tahoe-derived inferred profiles were calibrated into PANACEA expression space with *limma* ^51^. Matched controls were defined as the mean expression profiles of measured no-perturbation samples from the same cell line. The calibration model was fitted using matched cell-line control centroids only and did not use PANACEA drug-treatment conditions as calibration inputs. The resulting calibrated treatment profiles were used without subsequent clipping. Profile-concordance diagnostics compared calibrated and unaligned inferred profiles with measured PANACEA profiles, but measured PANACEA treatment profiles were not used to define the pathway-effect neighbour ranks.

#### Pathway-effect representation and drug neighbourhoods

Hallmark GSVA scores were computed for calibrated inferred treatment profiles and matched measured PANACEA control centroids using the same gene sets for all conditions ^48,52^. The PANACEA response-programme vector was defined as the treatment GSVA score vector minus the GSVA score vector of the matched measured PANACEA cell-line control centroid. No additional pathway-score standardisation was applied before ranking. Thus, the neighbourhood analysis used matched-control-relative pathway effects rather than absolute expression or uncentred pathway activity.

Drug neighbourhoods were ranked by Pearson correlation between pathway-effect vectors. For each query–candidate pair, similarities were computed within matched PANACEA cell-line contexts, and the drug-level similarity was the maximum similarity over the available cell-line contexts. Same-drug candidates were excluded from every candidate set, so top-1 concordance reflected a pharmacologically related compound rather than the query compound itself.

#### MoA annotation and top-1 concordance

Drug MoA annotations were obtained from the Broad Drug Repurposing Hub 2020-03-24 annotation table ^26^. These annotations were used only after ranking. Among the 32 PANACEA drugs, 29 had Broad Hub MoA annotations. The primary top-1 neighbour concordance analysis used the 27 drugs for which at least one candidate drug shared at least one MoA annotation. For query drug *i*, the top-1 concordance indicator was 1 if the highest-ranked candidate shared at least one Broad Hub MoA annotation with the query and 0 otherwise. The reported concordance was the mean of this indicator across the 27 evaluable query drugs.

Random-label nulls were computed by shuffling MoA label sets across drugs while keeping the inferred pathway-effect ranks and same-drug exclusion fixed. Label-prevalence expectations were also computed from the fraction of candidate drugs sharing at least one MoA annotation with each query drug. These query-specific probabilities defined the exact one-sided Poisson-binomial upper-tail test for the combined top-1 endpoint^27,28^. A Wilson 95% confidence interval was calculated for the observed top-1 success proportion. The first shared-MoA rank was the smallest candidate rank at which the query and candidate MoA sets intersected. Exact-identity occurrence subgroups, mean reciprocal rank, average precision and term-level recovery were secondary descriptive summaries.

### Colon development prediction

#### Dataset and split

Colon development experiments used a broad-stage conditioning variable. Following the gene-space scale used in published Squidiff examples, both models were evaluated on the same 800-HVG feature space. Enterocyte profiles were filtered, normalised to 10,000 counts per cell and log-transformed. The HVGs were selected on the complete dataset, so the feature-selection step was transductive. Post-conception week was encoded as early stage 1.0 for ≤ 14 pcw, transition stage 2.0 for 15–18 pcw and mature stage 3.0 for ≥ 19 pcw.

The model was trained on early and mature stages and evaluated on one pooled broad-stage-2 target containing the held-out PCW 16–18 cells. Broad stage 2 was supplied directly as the continuous conditioning value 2.0 and projected through the nonlinear conditioning MLP within the denoising network. No expression-space or latent-space midpoint of the early and mature cells was constructed.

#### Model training and inference

For colon development, the autoencoder used a 64-dimensional latent space, KL weight 2 × 10*^−^*^4^, dropout 0.2, batch size 256, learning rate 1 × 10*^−^*^4^, weight decay 3 × 10*^−^*^4^ and 15,000 update steps. The diffusion denoiser used four residual blocks, hidden dimension 256, context dimension 256, timestep embedding dimension 32 and dropout 0.2. It was trained with batch size 256, learning rate 1.5 × 10*^−^*^4^, weight decay 3 × 10*^−^*^4^, 1,200 training epochs and *v*-prediction. Inference generated held-out transition-stage profiles from the broad-stage conditioning variable, decoded predictions were clamped at zero and intermediate latent states were retained for reverse-diffusion process visualisation.

#### Colon development evaluation metrics

Colon prediction was evaluated on the pooled held-out PCW 16–18 enterocyte cells. The model generated all held-out cells with the single broad-stage-2 condition. Pseudo-bulk metrics were computed for the pooled transition target. PCW 16, 17 and 18 summaries were descriptive strata defined from cell annotations after generation and were not separate conditioning targets. Predicted and observed mean-expression vectors were compared over the shared 800-gene feature space.

UMAP visualisations used a fixed embedding for the combined ground-truth and generated cells. The embedding was computed from 50 principal components with 15 neighbours, minimum distance 0.3 and random seed 7. The same saved embedding positions were used for stage overlays and gene-expression overlays. Stage-feature trajectories used biology-guided epithelial genes selected from the 800-gene prediction space. Selection required paper-supported annotation, measurable held-out stage-2 expression, an interpretable early–transition–late pattern and at least one recovery criterion based on Wasserstein distance, mean-expression error or PCW-wise pseudo-bulk error. Trajectory panels report stage-wise mean log-normalised expression with standard error of the mean.

Single-cell distribution panels include all held-out PCW 16–18 cells, including zero-expression cells, and summarise gene-level expression-density recovery for representative developmental features. Pseudo-bulk error distributions and supplementary empirical cumulative distributions were computed as per-gene absolute differences between predicted and observed mean expression across the same 800-gene evaluation space.

Transition-stage epithelial composition was assessed using two exploratory modules guided by published fetal intestinal epithelial programmes^24^: an absorptive-trajectory module (*SELENOP*, *TTR*, *APOA1*, *APOB*, *FABP2*, *MUC13*, *GSTA1*, *RBP2*, *ALDOB*, *CUBN* and *MAMDC4*) and a BEST4/OTOP2-like module (*CA7*, *CA4*, *BEST4*, *OTOP2*, *GUCA2B*, *GUCA2A*, *KRT20*, *TFF3*, *LCN15*, *SPINK1* and *CEACAM5*). For each gene, the mean and standard deviation of log-normalised expression were estimated from observed cells across broad stages 1–3. Expression in the observed transition population and the PerturbLDM and Squidiff predictions was standardised using these fixed observed-reference parameters. The score for each module was the mean standardised expression of its constituent genes, and the programme margin was defined as the BEST4/OTOP2-like score minus the absorptive-trajectory score. Cells with a margin of at least 0.12 were assigned to the BEST4/OTOP2-like programme, those with a margin of at most −0.12 were assigned to the absorptive-trajectory programme, and the remaining cells were classified as mixed or low margin. Figure 4h reports the descriptive proportions obtained by applying these parameters and assignment rule unchanged to all three profiles.

### PBMC stimulation transfer

#### Dataset and split

PBMC stimulation experiments used the same architecture with categorical conditioning. Cells were filtered, normalised to 10,000 counts per cell and log-transformed. The Fig. 5 comparison used 2,000 highly variable genes selected on the complete dataset, so held-out stimulated cells informed the feature space. Each condition was defined by lineage and stimulation status. Stimulated B cells, CD8 T cells and FCGR3A^+^ monocytes were held out, while controls from those lineages remained in training. Evaluation comprised three held-out lineage tasks. Donors were not treated as independent inference units, although the dataset contained eight donors. This split evaluated stimulation-response transfer with retained control-state context rather than generalisation to lineages lacking such context.

#### Model training and inference

For PBMC, the autoencoder used a 128-dimensional latent space, KL weight 2 × 10*^−^*^4^, dropout 0.2, batch size 256, learning rate 1 × 10*^−^*^4^, weight decay 3 × 10*^−^*^4^ and 20,000 update steps. The diffusion denoiser used four residual blocks, hidden dimension *h* = 256, context dimension 256, timestep embedding dimension 32 and dropout 0.2. It was trained with batch size 256, learning rate 2 × 10*^−^*^4^, weight decay 3 × 10*^−^*^4^, 2,000 training epochs and *v*-prediction. Standard inference generated predictions for held-out stimulated cells, and decoded predictions were clamped at zero before lineage-level evaluation. Control-initialised predictions were generated across strengths from 0 to 1. Strength 0.4 was selected retrospectively by scoring the same held-out targets, and the resulting analysis was treated as a sensitivity analysis rather than an independently tuned comparator. Selected reverse-diffusion states are shown only as qualitative process diagnostics (Supplementary Figs. S11 and S12).

#### Expression and DEG evaluation

PBMC absolute-expression metrics were computed between predicted and measured stimulated mean-expression profiles for the same held-out cell type. Differential-expression analyses were matched-control relative. For each held-out cell type, the ground-truth DEG set was defined by comparing measured stimulated cells with measured control cells from the same cell type. Predicted DEG sets were defined by comparing each method’s predicted stimulated cells with the same matched control cells.

DEG-overlap metrics and the volcano/overlap display used bidirectional calls with adjusted *P <* 0.05, | log_2_ FC| *>* 0.25 and detection in at least 10% of either stimulated or control cells. Precision, recall, specificity and F1 were evaluated over the common post-detection gene universe, with *F* 1 = 2*PR/*(*P* + *R*). Adjusted *P* values were used to define within-profile feature sets rather than model-level inference because the analysis did not account for cells nested within donors. Highlighted response genes were selected from the most significant upregulated and downregulated ground-truth DEGs in the displayed FCGR3A^+^ monocyte comparison.

#### GO Biological Process analyses

For Fig. 5g and Supplementary Fig. S10, the enrichment analysis recomputed upregulated-gene lists using tie correct=True, adjusted *P <* 0.05, log_2_ FC *>* 0.25 and detection in at least 10% of stimulated cells. These enrichment inputs differed from the bidirectional DEG-overlap sets used for Fig. 5c–e. The focused display used seven terms selected from ground-truth enrichment results and listed in Supplementary Table S3. Gene membership was taken from the Enrichr GO Biological Process 2023 library and recorded in the companion source table. Term recovery was counted when a term was enriched at adjusted *P <* 0.05, and enrichment strength was summarised as − log_10_ adjusted *P*. These within-profile enrichment statistics were not interpreted as between-model significance tests.

GO-profile similarity was summarised by top-20 Jaccard overlap and by Spearman correlation of − log_10_ adjusted-*P* profiles.

Per-cell pathway scores were computed for the MSigDB Hallmark oxidative-phosphorylation (OXPHOS) gene set and the GO Biological Process fatty-acid beta-oxidation (FAO) set (GO:0006635)^48^. These sets contained 31 and 7 genes, respectively, in the common 2,000-HVG feature space. For each profile, a pathway score was defined as the mean expression of member genes minus the mean expression of expression-matched reference genes selected with 25 expression bins, 50 reference genes per bin and random seed 20260806. The FAO–OXPHOS composite was the arithmetic mean of the independently computed FAO and OXPHOS scores, giving equal weight to the two programmes. A lineage-level effect was calculated as the mean score of measured or predicted stimulated cells minus the mean score of measured lineage-matched control cells. For the effect comparison, we calculated the absolute deviation of each predicted composite effect from the measured composite effect within each lineage. Fig. 5h reports PerturbLDM error divided by scGen error for this comparison and for the corresponding one-dimensional Wasserstein distances between predicted and measured per-cell composite-score distributions. Values below 1 favour PerturbLDM. Supplementary Fig. S13 reports the separate FAO and OXPHOS effects. Its reference-mapped display used 30 principal components and a UMAP with 15 neighbours, minimum distance 0.3 and random seed 20260806 fitted only to the measured stimulated profiles. PerturbLDM and scGen profiles were transformed through the same PCA and UMAP models, and all panels used a common OXPHOS-score colour scale. These scores quantify expression of FAO- and OXPHOS-associated gene programmes rather than reaction flux or metabolite turnover. For complementary flux-informed analysis, HUMESS combines transcriptomic fold changes with genome-scale metabolic modelling to prioritise condition-specific reactions and associated genes ^53^.

### Evaluation metrics and statistical analysis

Expression-profile metrics comprised mean absolute error, mean squared error, root mean squared error, Pearson correlation, Spearman correlation, *R*^2^ and Chatterjee’s rank correlation ^54^. Matched-control-relative effect metrics comprised Pearson correlation, Spearman correlation and mean absolute error. For *R*^2^ and Chatterjee’s coefficient, observed profiles were supplied as the first argument and predicted profiles as the second. For condition-level comparisons, metrics were computed between predicted and observed mean profiles for the same held-out condition unless otherwise stated. In the nine primary Tahoe effect, signed-programme and distribution comparisons, the held-out condition was the paired unit (*n* = 13,942). We reported the median direction-oriented paired difference, a percentile 95% confidence interval from 2,000 condition-level bootstrap resamples (seed 20260718), the favourable-condition fraction and a two-sided paired Wilcoxon signed-rank test. Benjamini–Hochberg correction was applied separately to the effect family (three tests), signed-programme family (two tests) and distribution family (four tests). These intervals and tests quantify variation across the held-out conditions, not across independently trained models or independent atlases. Complete T01–T09 values are reported in Supplementary Table S4, and analysis units and denominators are summarised in Supplementary Table S5. Boxplots show interquartile ranges, centre lines show medians and whiskers show 5th–95th percentiles unless otherwise specified.

### Implementation

PerturbLDM was implemented in PyTorch, with DDPM scheduling from diffusers. Single-cell preprocessing used Scanpy and AnnData. Autoencoder and diffusion optimisation used AdamW, linear learning-rate decay and gradient clipping. Stochasticity arose from mini-batch sampling, VAE reparameterisation and diffusion sampling.

## Acknowledgements

The authors acknowledge the High-Performance Computing for Research facility at the Center for Secure Artificial Intelligence for Healthcare, McWilliams School of Biomedical Informatics, The University of Texas Health Science Center at Houston.

## Data availability

All primary datasets analysed in this study were previously published and are publicly available. Tahoe-100M is available from https://huggingface.co/datasets/tahoebio/Tahoe-100M. PANACEA gene-expression data are available from the Gene Expression Omnibus under accession GSE186341 and from Synapse at https://doi.org/10.7303/syn20968331. Human fetal-intestinal single-cell RNA-sequencing data are available under GEO accession GSE158702 and from Mendeley Data at https://doi.org/10.17632/gncg57p5x9.2. The PBMC data are available under GEO accession GSE96583. Derived data supporting the reported analyses are available from the corresponding author upon reasonable request.

## Code availability

PerturbLDM source code, documented benchmark workflows, configuration files and a compact PBMC example are publicly available under the MIT License at https://github.com/davidroad/PerturbLDM.

