## supplementary file for "PerturbLDM: conditional latent diffusion for modelling single-cell perturbation responses"

#### Supplementary Methods

Supplementary Figs. S1 and S2 summarise the PerturbLDM architecture, training behaviour and sensitivity to model configuration.

##### Tahoe random-split benchmark methods

**Evaluation unit, split and target definition.** The Tahoe random-split benchmark evaluated 46,471 measured treatment conditions, each defined by one drug, dose and cell line: 32,529 formed the development set and 13,942 were retained for held-out evaluation. Within the development set, a fixed 10% condition-level validation subset was generated with random seed 42, yielding 29,277 fitting and 3,252 validation conditions. This partition was shared by MLP, random forest, CPA and chemCPA, with no condition overlap among the fitting, validation and held-out evaluation sets. All 379 drugs, 47 cell lines and three dose levels (0.05, 0.5 and 5.0 $\mu$M) occurred in both the development and held-out evaluation sets, so the benchmark evaluates unmeasured combinations of observed factors rather than unseen drugs, cell lines or dose levels. For the supervised MLP and random-forest baselines, each condition was represented by the mean expression vector over its constituent cells. A matched-control reference vector was computed separately for each cell line as the mean expression vector over control cells from that cell line. The supervised learning target was the matched-control-relative response,

$$\Delta\mathbf{x}_{c,d,\mu} = \bar{\mathbf{x}}_{c,d,\mu} - \bar{\mathbf{x}}_{c}^{\mathrm{ctrl}},$$

where $c$ indexes cell line, $d$ indexes drug and $\mu$ indexes dose. Predictions were converted back to absolute expression before evaluation as

$$\hat{\mathbf{x}}_{c,d,\mu} = \bar{\mathbf{x}}_{c}^{\mathrm{ctrl}} + \widehat{\Delta\mathbf{x}}_{c,d,\mu}.$$

This formulation trains the supervised baselines to model the perturbation-induced deviation while all reported expression metrics compare predicted and observed profiles on the same absolute expression scale. The zero-effect control predicted $\widehat{\Delta\mathbf{x}} = 0$ for every condition and therefore corresponds to using the matched cell-line control mean as the predicted treated profile.

**Supervised baseline variable construction.** For the supervised baselines, condition-level design matrices combined drug identity, cell-line identity, dose and the matched-control expression profile for the target cell line. Drug and cell-line one-hot encoders and the dose standardisation were fitted using the 29,277 fitting conditions and then applied unchanged to the validation and held-out evaluation conditions. The MLP received 379 drug indicators, 47 cell-line indicators, standardised dose and the full 13,784-gene matched-control profile. The random forest received the same encoded condition variables and 1,500 matched-control genes selected by weighted variance across cell-line control profiles using fitting conditions only. No response expression from the validation or held-out evaluation conditions contributed to variable construction.

**MLP baseline.** The MLP baseline was a multi-output feed-forward regressor trained to predict the full $\Delta$-expression vector. Its architecture was 14,211 $\rightarrow$ 512 $\rightarrow$ 256 $\rightarrow$ 13,784, with batch normalisation and ReLU after the first hidden layer, ReLU after the second hidden layer and a linear output. A compact sequential search compared learning rates $10^{-4}$, $5 \times 10^{-4}$ and $10^{-3}$, followed by dropout values 0, 0.1, 0.2 and 0.3 at the selected learning rate. Selection used pooled validation $\Delta$-expression mean-squared error. The selected model used Adam, learning rate $10^{-4}$, batch size 1,024, no dropout and no weight decay. It was trained for 100 epochs, and the epoch-99 model with the lowest validation loss was used to predict the held-out responses. The held-out evaluation responses were not used for hyperparameter or epoch selection.

**Random-forest baseline.** The random-forest baseline was a multi-output $\Delta$-expression regressor with 1,927 inputs comprising encoded drug, cell-line and dose variables and 1,500 matched-control genes selected from the fitting conditions. The model used 300 trees, a maximum depth of 10, square-root variable sampling, bootstrap sampling and random seed 42. Minimum leaf sizes of 5, 20 and 50 were compared using validation $\Delta$-expression mean-squared error, which selected a minimum leaf size of 5. Held-out evaluation responses were excluded from model selection.

**CPA baseline.** CPA used the shared fitting, validation and held-out evaluation partitions, with drug identity as the perturbation variable, dose as the dosage variable and cell line as the categorical covariate. Adversarial class weights were computed from fitting cells only, and `DMSO-TF` was used as the control group. The model used Gaussian reconstruction loss, global seed 0, batch size 131,072, learning rate $10^{-3}$, adversarial regularisation weight 200 and adversarial penalty weight 400. It was trained for up to 20 epochs, with validation after every epoch and early stopping on the CPA validation metric with patience 4 and minimum improvement $10^{-4}$. Early stopping ended training after eight epochs, and the epoch-4 model, selected by the highest validation metric, was used to generate predictions from the corresponding `DMSO-TF` controls under the requested drug-dose perturbations.

**chemCPA baseline.** chemCPA used the same fitting, validation and held-out evaluation partitions and matched controls as the other learned comparators. Drug structure was represented by a combined SMILES feature encoding containing 1,024-bit Morgan fingerprints with radius 2 and 300 RDKit descriptors, with feature standardisation. Drug identity was used as the perturbation variable, dose as the dosage variable, cell line as a categorical covariate and `DMSO-TF` as the control group.

The chemCPA implementation used a ComPert-style architecture with SMILES-aware inputs. It used a latent dimension of 1,024, encoder and decoder widths of 2,048, and depths of 4 for the encoder, decoder and autoencoder. The adversary had width 1,024 and depth 3, the embedding encoder width 1,024 and depth 3, and the dose encoder width 512 and depth 3. The decoder used a linear activation and a log-sigmoid dose response. The selected run used five epochs, batch size 2,048, learning rate $10^{-3}$, weight decay $10^{-5}$, Adam optimisation, a ReduceLROnPlateau scheduler and validation-loss early stopping with patience 3. The model with the lowest validation loss was used to generate predictions with the integrated SMILES features. Supplementary Fig. S3e–h summarises training and model selection for each comparator. Numerical loss values are not comparable across models, and random forest has no epochwise loss trace.

**Benchmark evaluation and statistical comparison.** For each held-out condition, absolute-expression metrics were computed by comparing the predicted and observed gene-expression vectors. Distribution metrics were computed for PerturbLDM, CPA and chemCPA, which generated condition-aligned single-cell samples. The supervised mean-profile baselines were evaluated at the profile, effect and programme levels. For the nine Tahoe comparisons (T01–T09), the held-out condition was the paired unit ($n$ = 13,942). We reported median paired differences oriented so that positive values favoured PerturbLDM, together with percentile 95% confidence intervals from 2,000 condition bootstrap resamples (seed 20260718), favourable-condition fractions and two-sided asymptotic paired Wilcoxon signed-rank tests. Zero differences were excluded, and no continuity correction was applied. Benjamini–Hochberg adjustment was applied separately to the three effect, two signed-programme and four distribution endpoints. Complete values are reported in Supplementary Table S4, and analysis units and denominators are summarised in Supplementary Table S5.

#### Study-specific benchmark methods

##### scGen for PBMC stimulation transfer

The Fig. 5 comparison used the same 2,000 highly variable genes (HVGs) and fitting and held-out evaluation split as the PerturbLDM analysis. These genes were selected using the complete dataset, including held-out cells. The comparison comprised three held-out lineage-specific stimulated states without donor-level replication or replicate model fits. The fitting set contained 11,842 cells, and the held-out evaluation set contained 1,710 stimulated cells. Stimulated B cells, CD8 T cells and FCGR3A$^+$ monocytes were withheld from training while their control-state cells were retained. CD4 T cells, CD14$^+$ monocytes, dendritic cells and NK cells provided training cell types with both control and stimulated states.

The scGen variational autoencoder was trained for up to 100 epochs with batch size 32 and early-stopping patience 25. Training stopped after 26 epochs. For prediction, a single global stimulation vector was estimated in scGen latent space from training cell types only. Control and stimulated training cells were balanced across cell-type labels by sampling equal numbers of control and stimulated cells without replacement. The perturbation vector was computed as

$$\Delta \mathbf{z}_{\mathrm{IFN}} = \overline{\mathbf{z}}_{\mathrm{stim,train}} - \overline{\mathbf{z}}_{\mathrm{ctrl,train}}.$$

For each held-out cell type $k$, control-state cells from that cell type were encoded to latent vectors, shifted by $\Delta \mathbf{z}_{\mathrm{IFN}}$, and decoded through the scGen generative decoder.

$$\widehat{\mathbf{x}}_{\mathrm{stim},k} = \mathrm{Dec}_{\mathrm{scGen}}\{\mathbf{z}_{\mathrm{ctrl},k} + \Delta \mathbf{z}_{\mathrm{IFN}}\}.$$

No held-out stimulated expression profiles were used to train scGen or to estimate the latent shift.

The concatenated scGen predictions for the three held-out cell types were evaluated with the same held-out stimulated profiles and matched-control profiles used for PerturbLDM. Absolute-expression metrics were computed on condition-level gene means over the common gene set and included Pearson correlation, Spearman correlation, $R^2$, RMSE and MAE. DEG overlap used the bidirectional Wilcoxon calls defined in the main Methods. For pathway recovery, upregulated-gene lists were recomputed before Enrichr analysis over GO Biological Process 2023, MSigDB Hallmark 2020 and KEGG 2021 Human. The focused PBMC pathway analysis evaluated recovery of the seven GO Biological Process terms listed in Supplementary Table S3, together with their enrichment strength and rank concordance with the measured-response enrichment results. The seven displayed GO terms were selected from the measured-response analysis by the authors rather than prespecified. Within-profile adjusted $P$ values were used to define gene-set calls, not to compare methods. FAO, OXPHOS and composite-score analyses used the same measured and predicted cells and the scoring procedure defined in the main Methods.

##### Squidiff for fetal-colon maturation prediction

Squidiff served as the study-specific comparator for fetal-colon maturation prediction. Early PCW 11–14 and mature PCW 19–23 enterocyte states provided observed anchors, and both methods generated one pooled broad-stage-2 population evaluated against 1,529 held-out PCW 16–18 cells. Squidiff generated this population using its semantic-latent interpolation procedure between the two anchors. Exact PCW was not a generation condition and was used only for descriptive stratification after generation. Following the gene-space scale used in published Squidiff examples, both models were evaluated over the same set of 800 HVGs, which was selected using the complete dataset, including held-out cells. Because gene selection used the complete dataset and the evaluation comprised one pooled target population, the comparison is descriptive and does not estimate variation across biological replicates.

#### Supplementary Figures

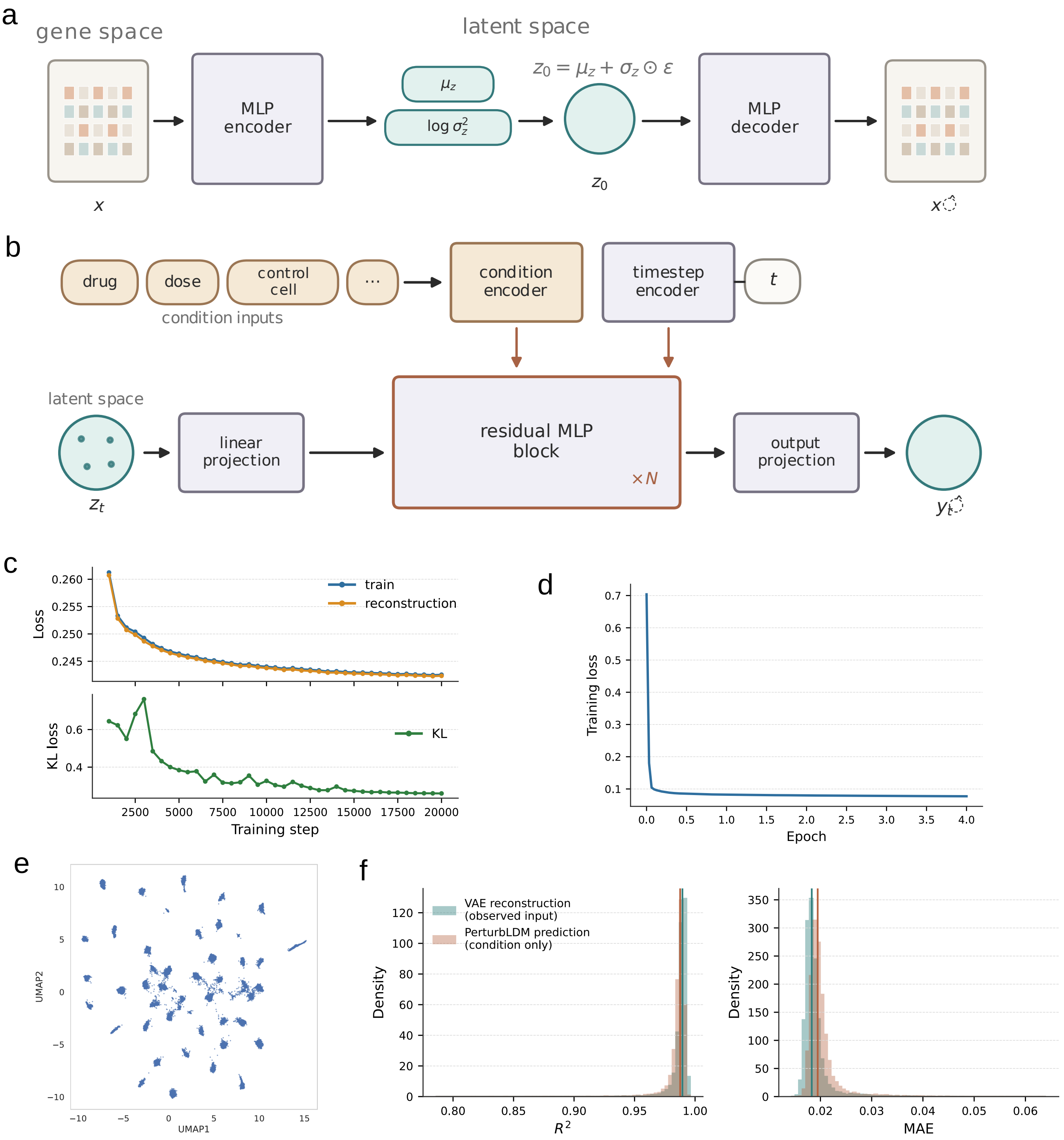


**Supplementary Fig. S1. PerturbLDM architecture and training behaviour. a,** Variational autoencoder architecture for mapping high-dimensional gene-expression profiles into a latent transcriptomic state space and decoding latent states back to gene space. **b,** Conditional denoiser schematic. Drug identity, dose and matched-control cell state are encoded together with the diffusion timestep and used by residual MLP blocks to predict the diffusion target in latent space. **c,** VAE reconstruction-loss and KL-loss traces across training. **d,** Diffusion-stage training-loss trace for the conditional denoiser. **e,** UMAP visualisation of encoded VAE latents, providing a qualitative view of latent organisation. **f,** Distributions of metrics for VAE reconstruction of observed profiles and PerturbLDM conditional generation of held-out responses. VAE reconstruction assesses the encode–decode representation, whereas PerturbLDM assesses condition-based generation. The panels characterise different stages of the framework and are not directly comparable.

**a**

| Variant | Drug embedding | Diffusion target | Latent dim. |
|---|---|---|---|
| **PerturbLDM** | Trainable | $v$ | 1024 |
| **Frozen SMILES** | Frozen SMILES drug emb. | $v$ | 1024 |
| $z_0$ **target** | Trainable | $z_0$ | 1024 |
| $\varepsilon$ **target** | Trainable | $\varepsilon$ | 1024 |
| **Latent 128** | Trainable | $v$ | 128 |

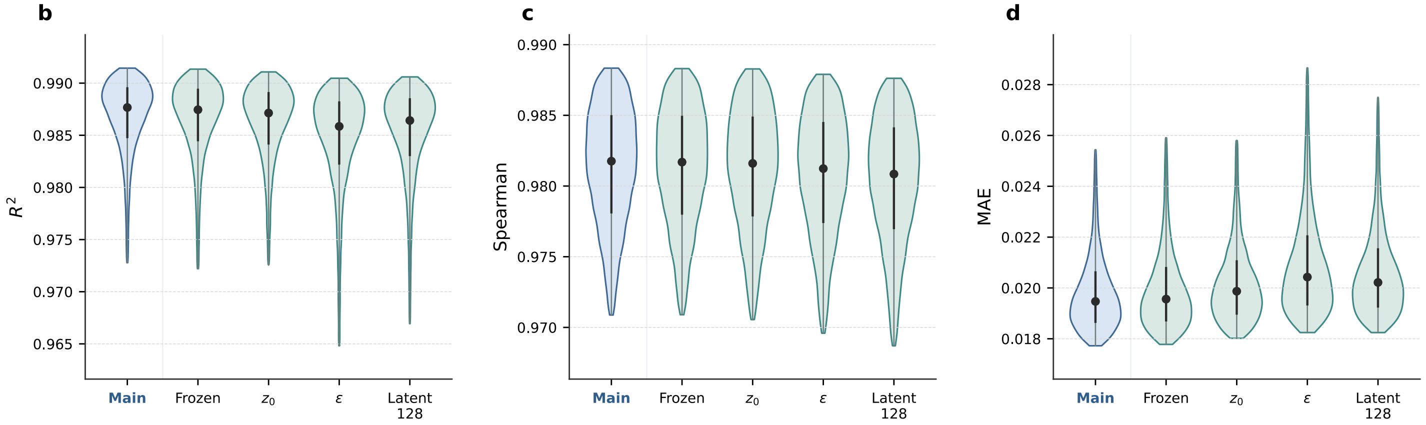


**Supplementary Fig. S2. Sensitivity to PerturbLDM model configurations. a,** PerturbLDM model configurations evaluated in the Tahoe setting, including the main configuration, frozen SMILES drug embeddings, alternative diffusion prediction targets and a reduced latent dimension. **b–d,** Distributions of condition-level $R^2$, Spearman correlation and MAE across variants. Each displayed variant contributes $n = 13{,}942$ aligned held-out conditions to each metric. The panels provide single-run sensitivity summaries across the evaluated configurations.

### Tahoe analyses

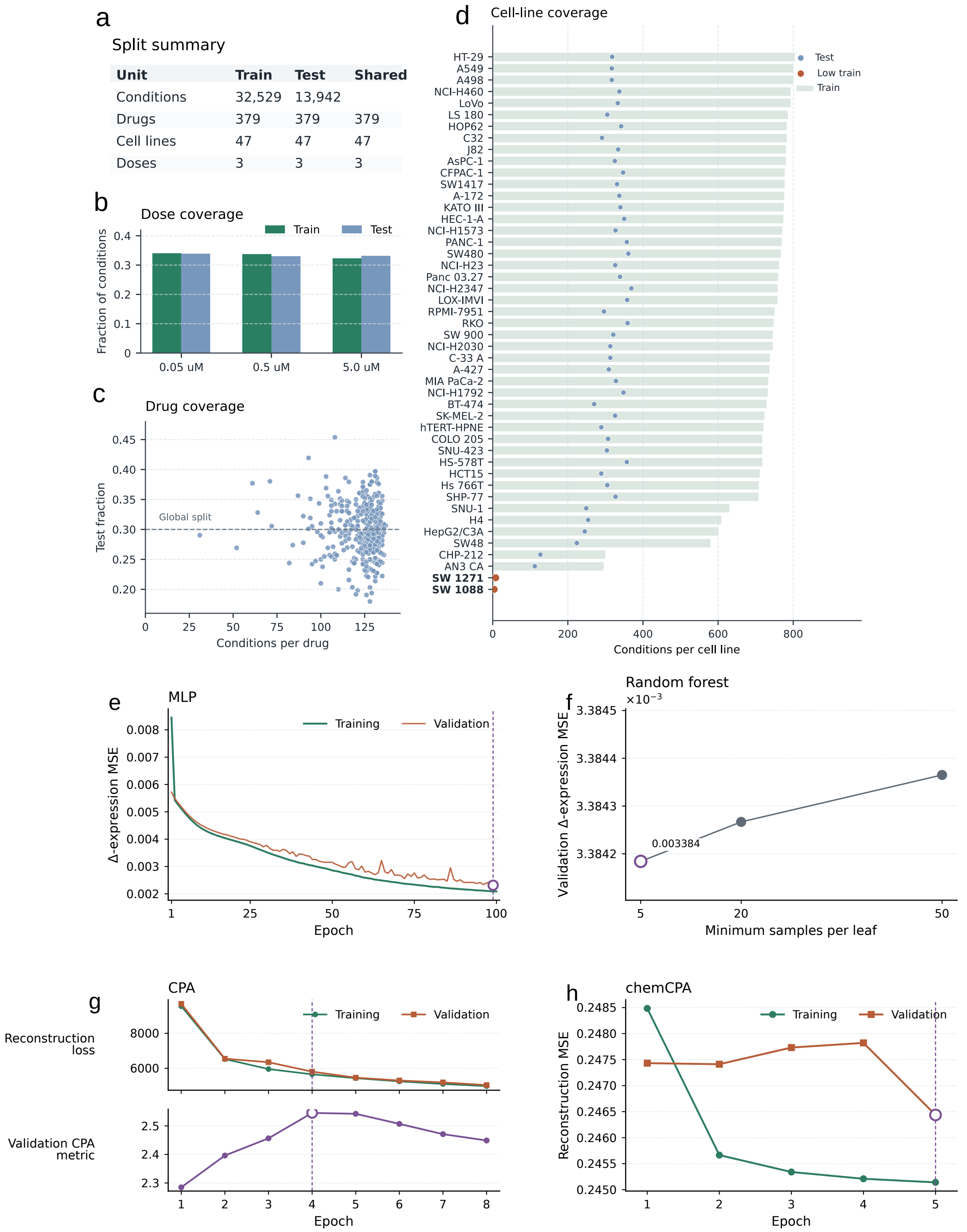


**Supplementary Fig. S3. Tahoe benchmark design and learned-comparator selection. a,** Development and held-out evaluation split for 46,471 analysed Tahoe treatment conditions, where each condition is defined by one drug, dose and cell line (32,529 development and 13,942 held-out evaluation conditions). All 379 drugs, 47 cell lines and three dose levels occurred in both splits, defining an observed-factor combination benchmark. **b,** Dose coverage in the development and held-out evaluation sets across the three assayed doses (0.05, 0.5 and 5.0 $\mu$M). **c,** Drug-level held-out fractions relative to the global split fraction, showing heterogeneous split coverage across drug groups. **d,** Cell-line coverage, with development-condition bars and held-out-evaluation-condition points for each cell line. Condition counts vary across cell lines. Coverage is not a calibrated estimate of prediction uncertainty. **e,** MLP training and validation matched-control-relative ($\Delta$-expression) mean-squared error across 100 epochs. The epoch-99 model, marked by the open circle, minimised validation error and was used to predict the held-out responses. **f,** Random-forest validation $\Delta$-expression mean-squared error for the three prespecified minimum-leaf sizes. A minimum leaf size of 5 gave the lowest validation error. The small separation among settings indicates limited sensitivity within this compact search. Random forest is fitted once per hyperparameter setting and therefore has no epochwise training curve. **g,** CPA training and validation reconstruction losses (top) and the CPA validation metric used for model selection (bottom). Validation was evaluated after every epoch. The validation metric was maximal at epoch 4. Training stopped after epoch 8 under the prespecified patience of four epochs. **h,** chemCPA training and validation reconstruction mean-squared error across five epochs. The epoch-5 model had the lowest validation reconstruction error. For all four learned comparators, the development set comprised 29,277 fitting and 3,252 validation conditions with identical condition membership. The held-out evaluation set of 13,942 conditions was excluded from model selection. Optimisation objectives and loss scales differ among methods, and these panels document model-specific selection rather than cross-method loss comparison or run-to-run variability.

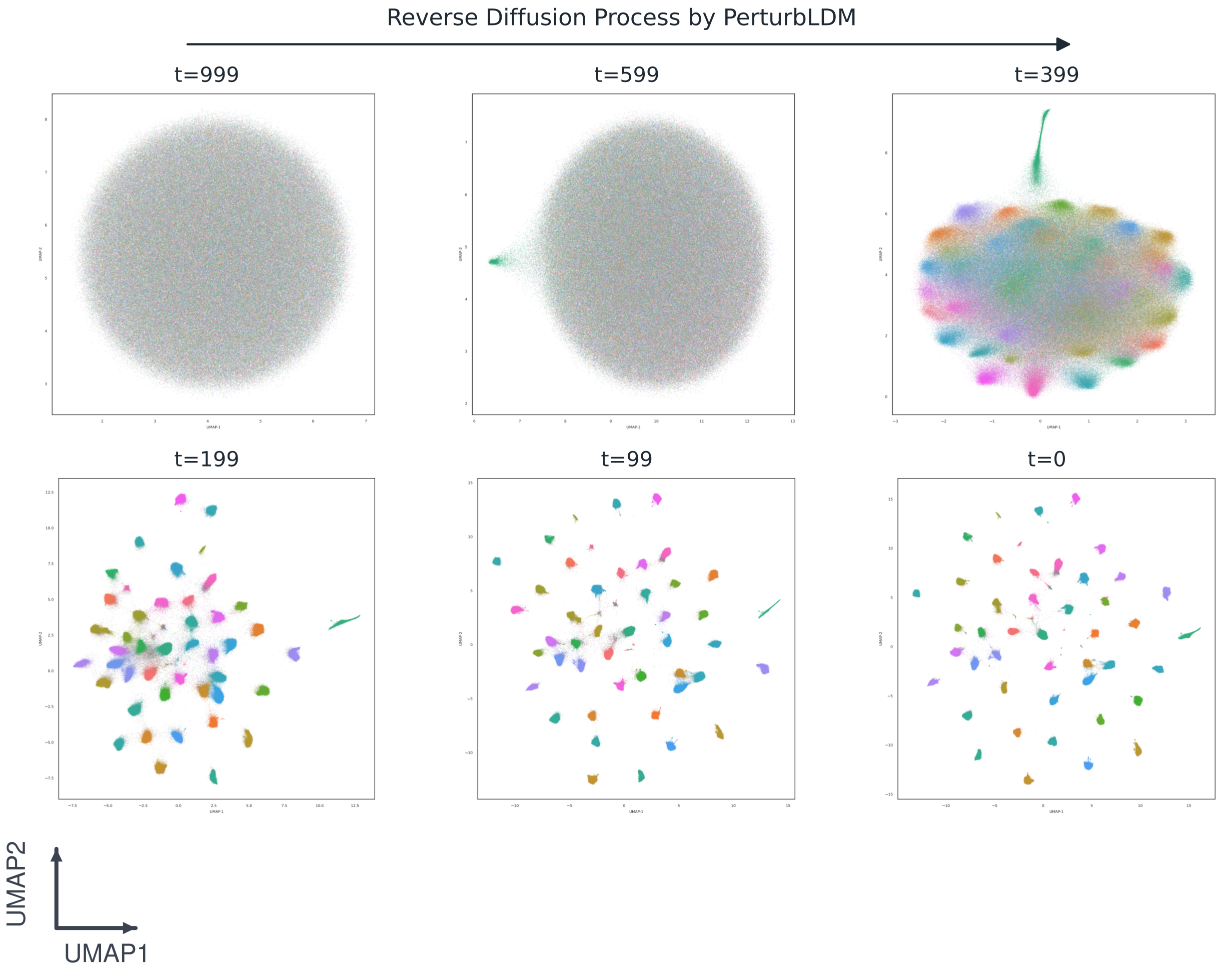


**Supplementary Fig. S4. Timestep-specific Tahoe reverse-diffusion latents.** UMAP visualisations show latent samples at selected reverse-diffusion timesteps, from high noise to the final latent state. Points are coloured by consistent anonymous cell-line categories. The UMAP axes indicate embedding orientation. Each timestep was embedded separately, so positions are not directly comparable and the panels do not quantify prediction accuracy.

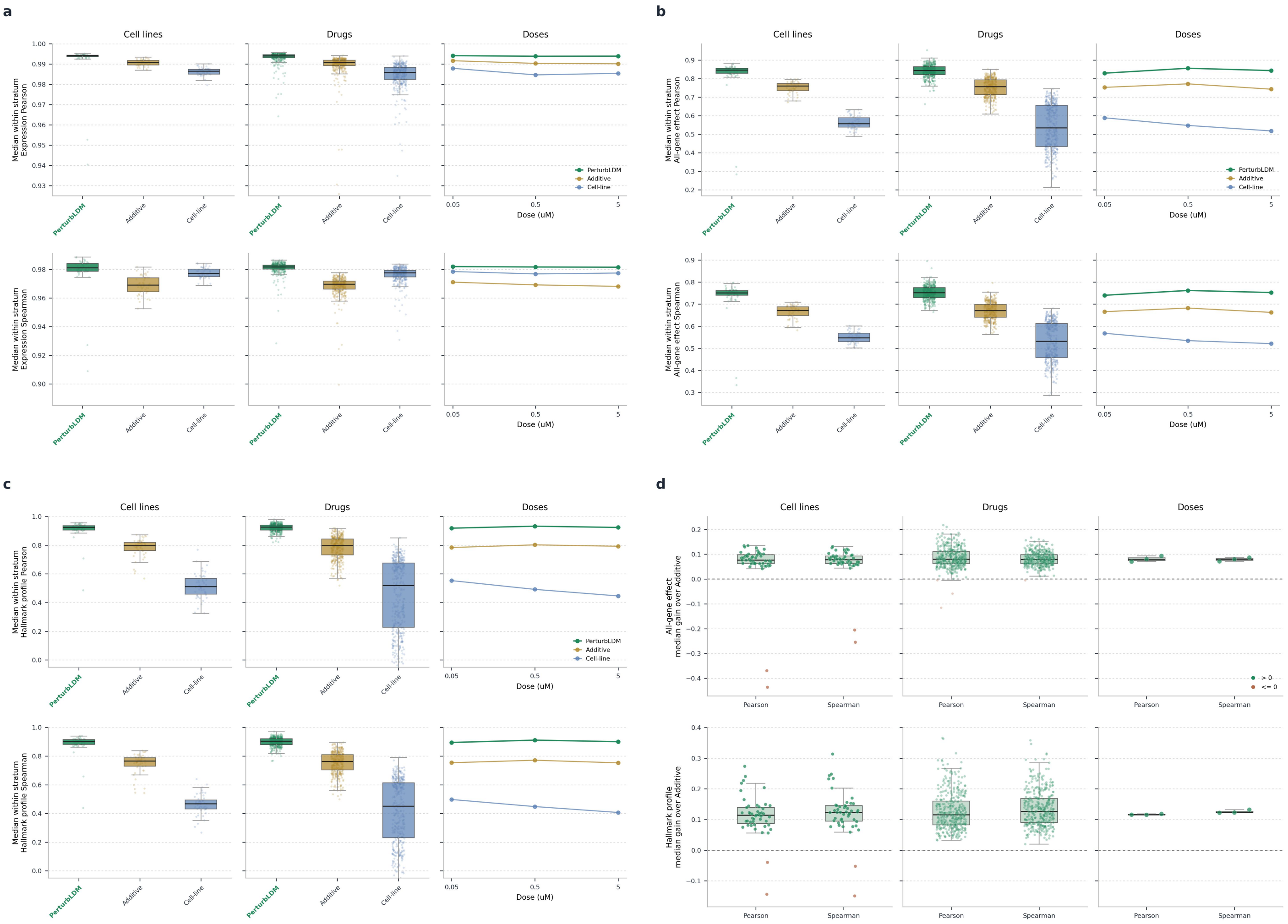


**Supplementary Fig. S5. Tahoe response concordance across drug, dose and cell-line strata. a,** Expression-level Pearson and Spearman metrics stratified by cell line, drug and dose for PerturbLDM, AdditiveMean and CellLineMean. **b,** Matched-control-relative all-gene effect Pearson and Spearman metrics stratified by the same biological axes. **c,** Signed Hallmark effect-profile Pearson and Spearman metrics stratified by cell line, drug and dose. Hallmark profiles are signed mean matched-control-relative effect summaries over Hallmark gene-set members, not enrichment, ssGSEA or GSVA scores. **d,** Paired gain over AdditiveMean for all-gene effect and Hallmark profile metrics. For panels **a–c**, each point summarises one stratum and represents the median of condition-level metrics among held-out conditions in that stratum. For panel **d**, gains are computed at the matched condition level as PerturbLDM minus AdditiveMean and then summarised within each stratum. Per-stratum condition counts and eligibility rules are supplied with the source data. Strata are descriptive summaries of the same held-out conditions, not independent replicates or inferential units.

### PANACEA pathway-neighbourhood analyses

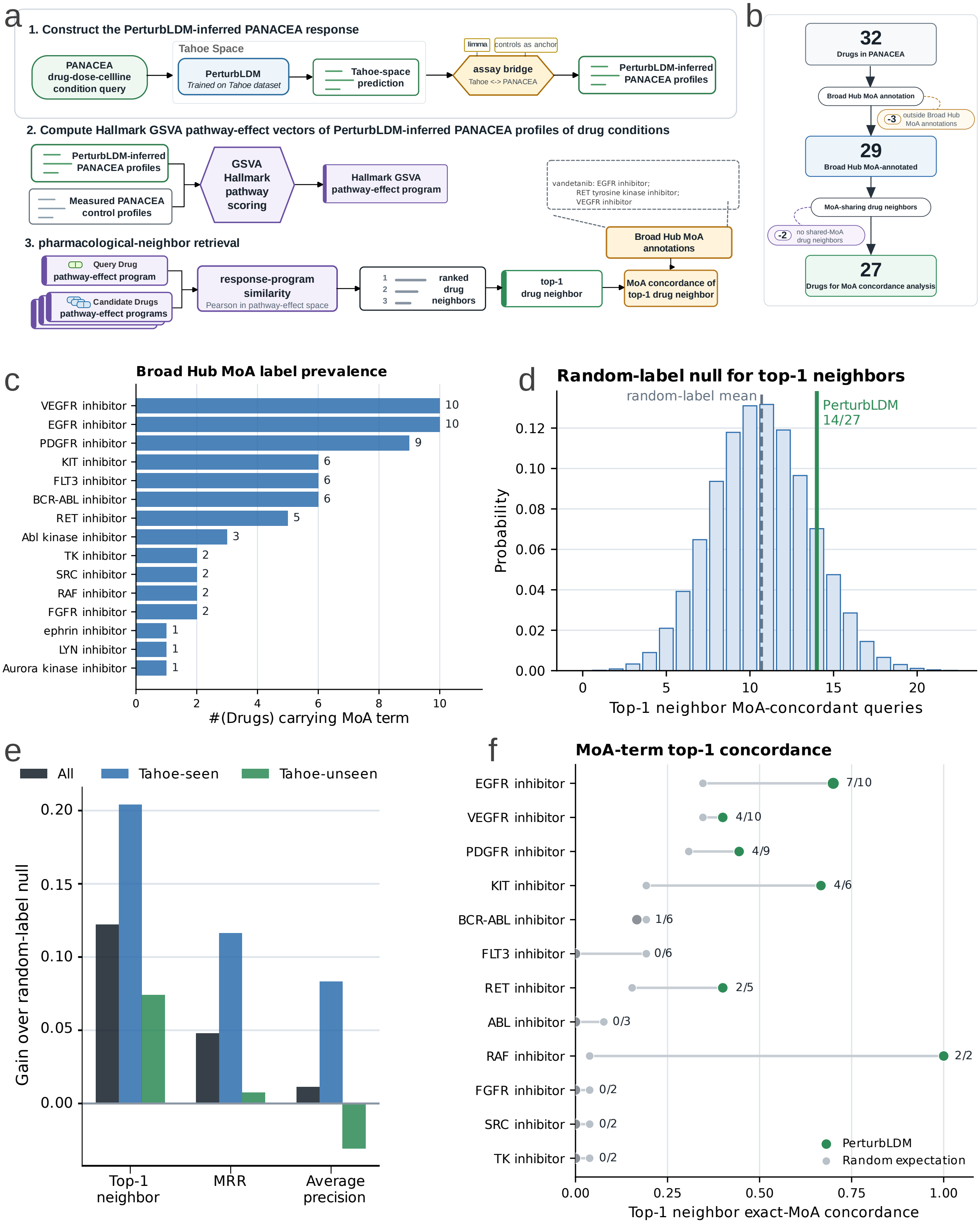


**Supplementary Fig. S6. PANACEA top-1 neighbour concordance and null analyses. a,** PANACEA response-programme neighbourhood workflow. Precomputed profiles generated by a Tahoe-trained frozen-SMILES variant are aligned to PANACEA space using matched cell-line controls, converted to matched-control-relative Hallmark GSVA pathway-effect vectors and ranked by similarity after same-drug exclusion. Neither measured

PANACEA treatment profiles nor MoA labels contributed to the ranking. MoA annotations were applied afterwards for evaluation.

**b,** Evaluation-set definition. The analysis starts from 32 PANACEA drugs. It retains 29 drugs with Broad Drug Repurposing Hub MoA annotations and evaluates 27 drugs with at least one MoA-sharing candidate.

**c,** Broad Hub MoA label prevalence in the 27-drug evaluation set. Bars indicate the number of drugs carrying each MoA annotation. These prevalences define the label-based expectations and random-label nulls used in the analysis.

**d,** Null analyses for top-1 neighbour concordance. This endpoint was observed for 14 of 27 drugs (51.9%, Wilson 95% CI, 34.0–69.3%). The exact query-specific Poisson-binomial test did not reach significance (one-sided $P = 0.1207$). A random-label permutation provided a secondary null comparison.

**e,** Gain over the corresponding random-label null for top-1 concordance, mean reciprocal rank and average precision, shown overall and by exact-identity-present or exact-identity-absent strata. The artwork's seen/unseen labels do not denote a prospective drug-held-out split.

**f,** Exact MoA-term top-1 recovery. Coloured points show observed term-specific concordance, grey points show the corresponding random expectation, and labels give the number of successful and eligible queries.

#### Fetal-colon maturation analyses

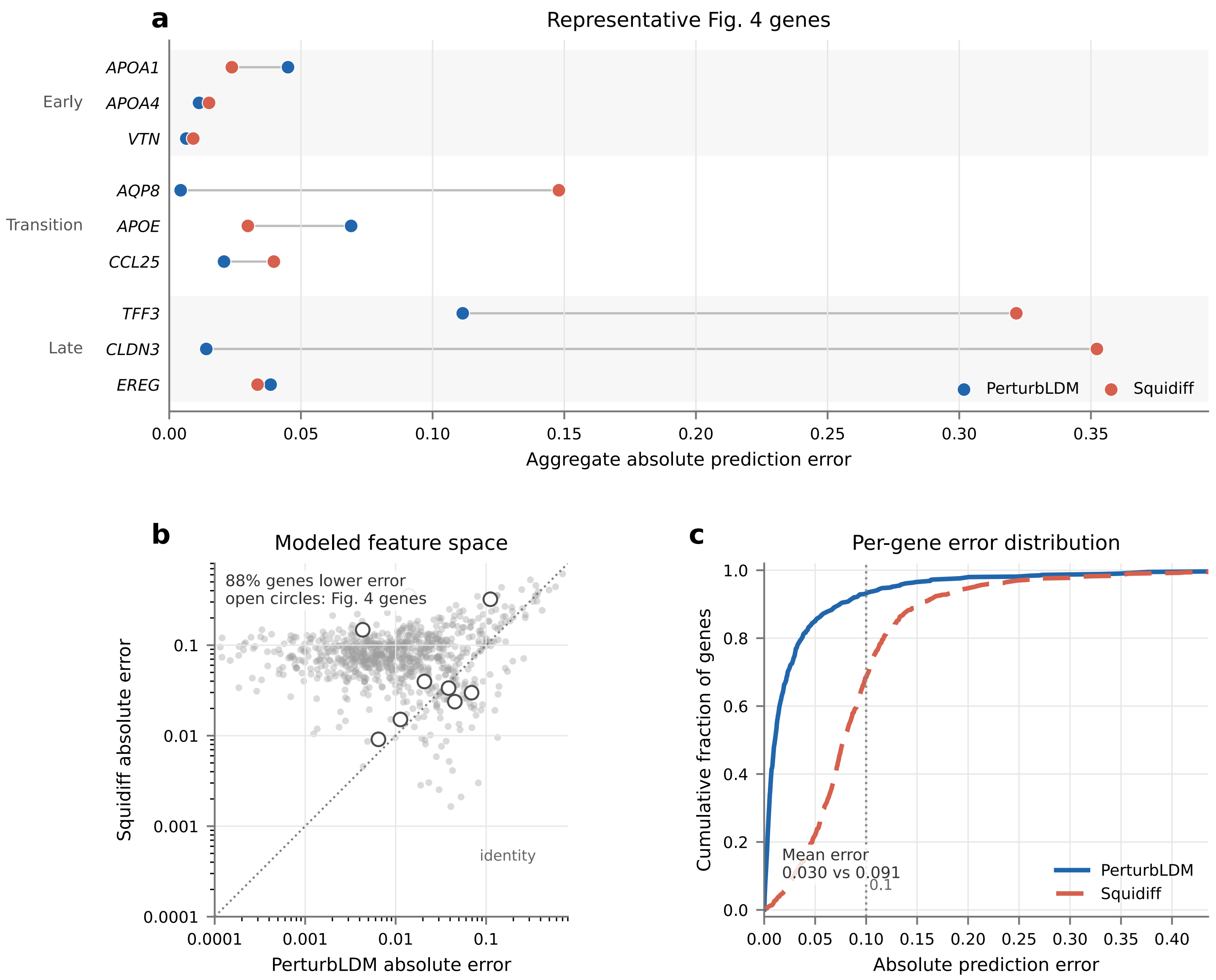


**Supplementary Fig. S7. Fetal-colon prediction errors across selected and shared genes. a,** Absolute prediction errors for the nine Fig. 4 genes selected retrospectively using published annotations and prespecified recovery criteria. **b,** PerturbLDM and Squidiff per-gene errors across the shared set of 800 genes. Points above the identity line indicate lower PerturbLDM error. **c,** ECDF of per-gene absolute errors. At a descriptive tolerance of 0.1 on the log-normalised expression scale, PerturbLDM predicts 745 of 800 genes (93.1%) within tolerance, compared with 547 of 800 (68.4%) for Squidiff. Genes are correlated measurements within one pooled developmental comparison, not biological replicates. The set of 800 HVGs was selected using the complete dataset.

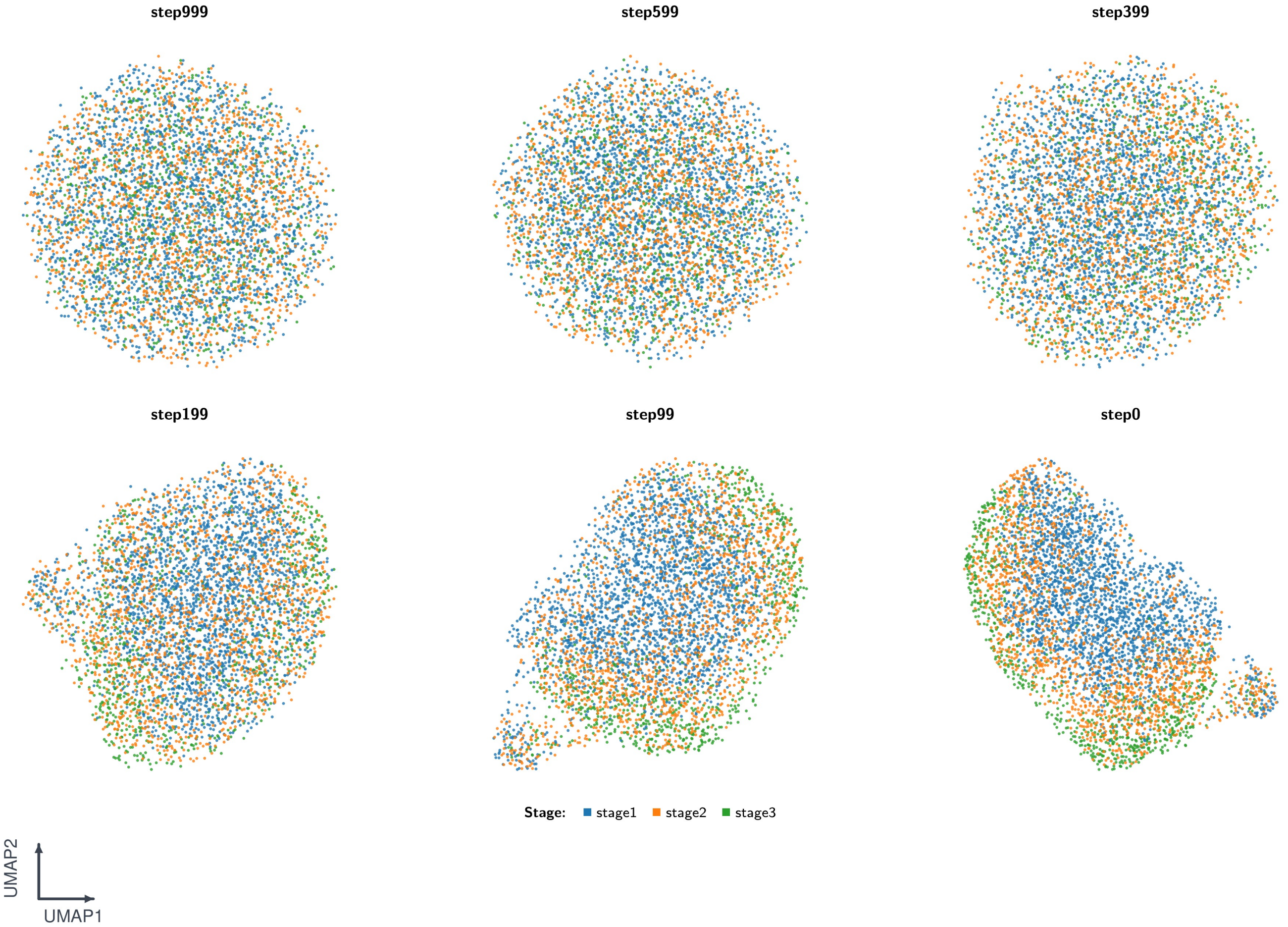


**Supplementary Fig. S8. Developmental reverse-diffusion latent states.** UMAP visualisations show selected PerturbLDM latent states for the Fig. 4 fetal-colon maturation analysis at step999, step599, step399, step199, step99 and step0. Panel labels report the post-update diffusion index, so the saved latent_1000 snapshot appears as step999. The legend follows the Fig. 4a stage colouring (stage1, stage2 and stage3), and a UMAP1/UMAP2 key indicates the orientation of each embedding. Each timestep was embedded separately, so cross-timestep positions are not directly comparable. These embeddings describe denoising progress and do not represent a biological developmental trajectory.

### PBMC analyses

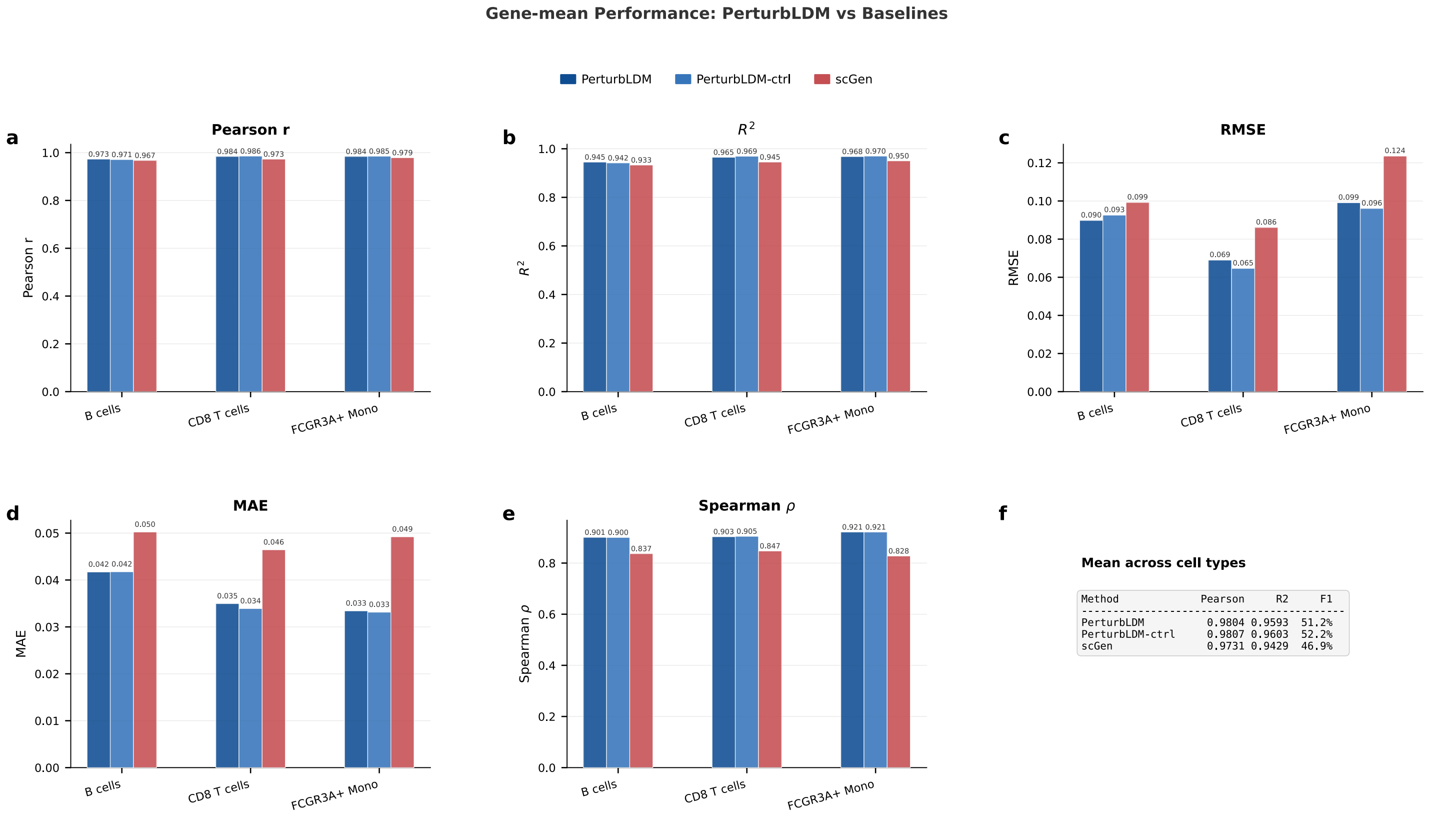


**Supplementary Fig. S9. PBMC expression-prediction metrics by held-out cell type.** Panels **a–e** show lineage-level gene-mean metrics for PerturbLDM, PerturbLDM-ctrl and scGen across held-out IFN-$\beta$-stimulated B cells, CD8 T cells and FCGR3A$^+$ monocytes: **a,** Pearson correlation, **b,** $R^2$, **c,** RMSE, **d,** MAE and **e,** Spearman correlation. Higher correlations and $R^2$, together with lower RMSE and MAE, indicate predictions closer to the measured stimulated profiles. **f,** Unweighted means across the three held-out lineages. Values are descriptive ($n = 3$ lineages), and PerturbLDM-ctrl used a strength selected retrospectively against the held-out targets. Exact lineage-level values are reported in Supplementary Table S6.

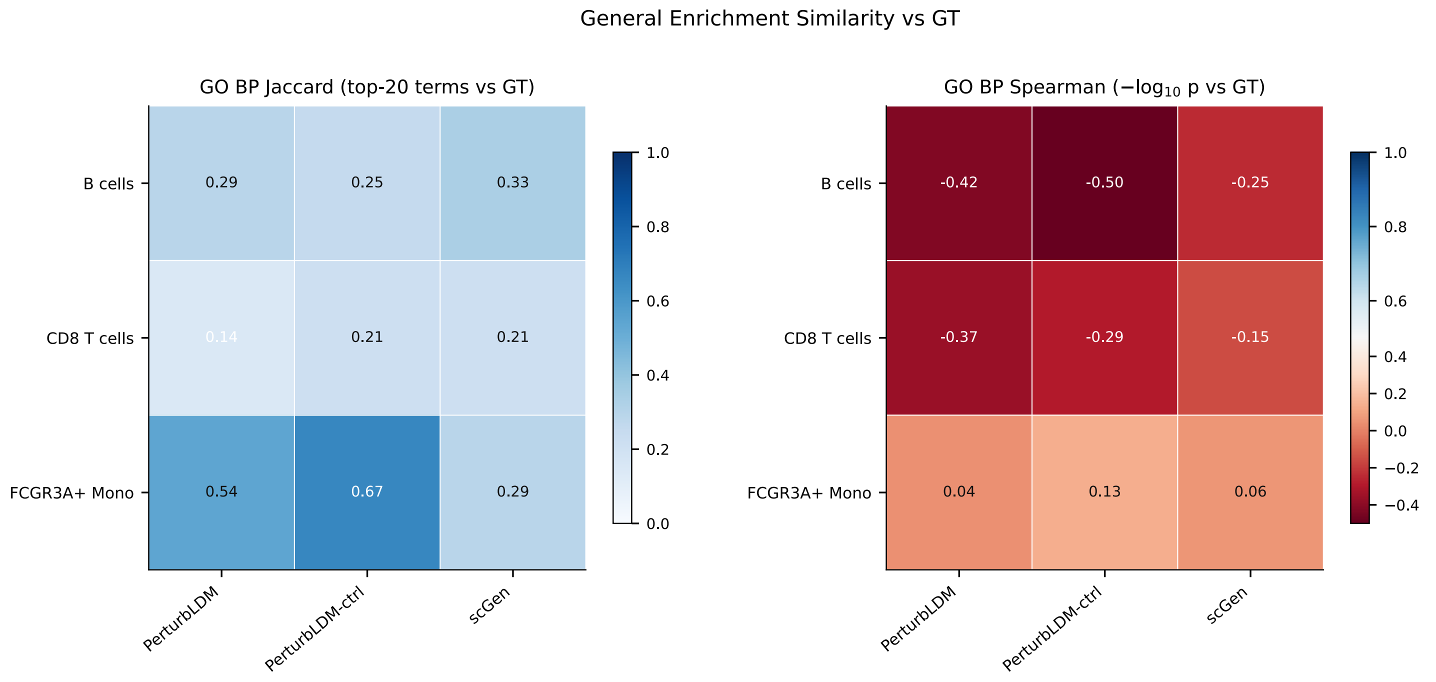


**Supplementary Fig. S10. PBMC GO enrichment similarity to measured responses.** Rows denote held-out B cells, CD8 T cells and FCGR3A$^+$ monocytes. Columns denote PerturbLDM, PerturbLDM-ctrl and scGen. Left, Jaccard overlap with the corresponding top-20 GO Biological Process terms from measured responses. Right, Spearman correlation of $-\log_{10}$ adjusted enrichment-$P$ profiles with the measured-response enrichment profile. Higher values indicate greater similarity to the measured enrichment profile. Enrichment profiles were computed from the separate upregulated-gene workflow defined in Methods, not from the bidirectional overlap sets used for Fig. 5e. The focused seven-term display in Fig. 5g uses author-selected terms from the measured-response analysis. Within-profile enrichment-adjusted $P$ values and numbers of GO terms do not constitute method-comparison sample sizes.

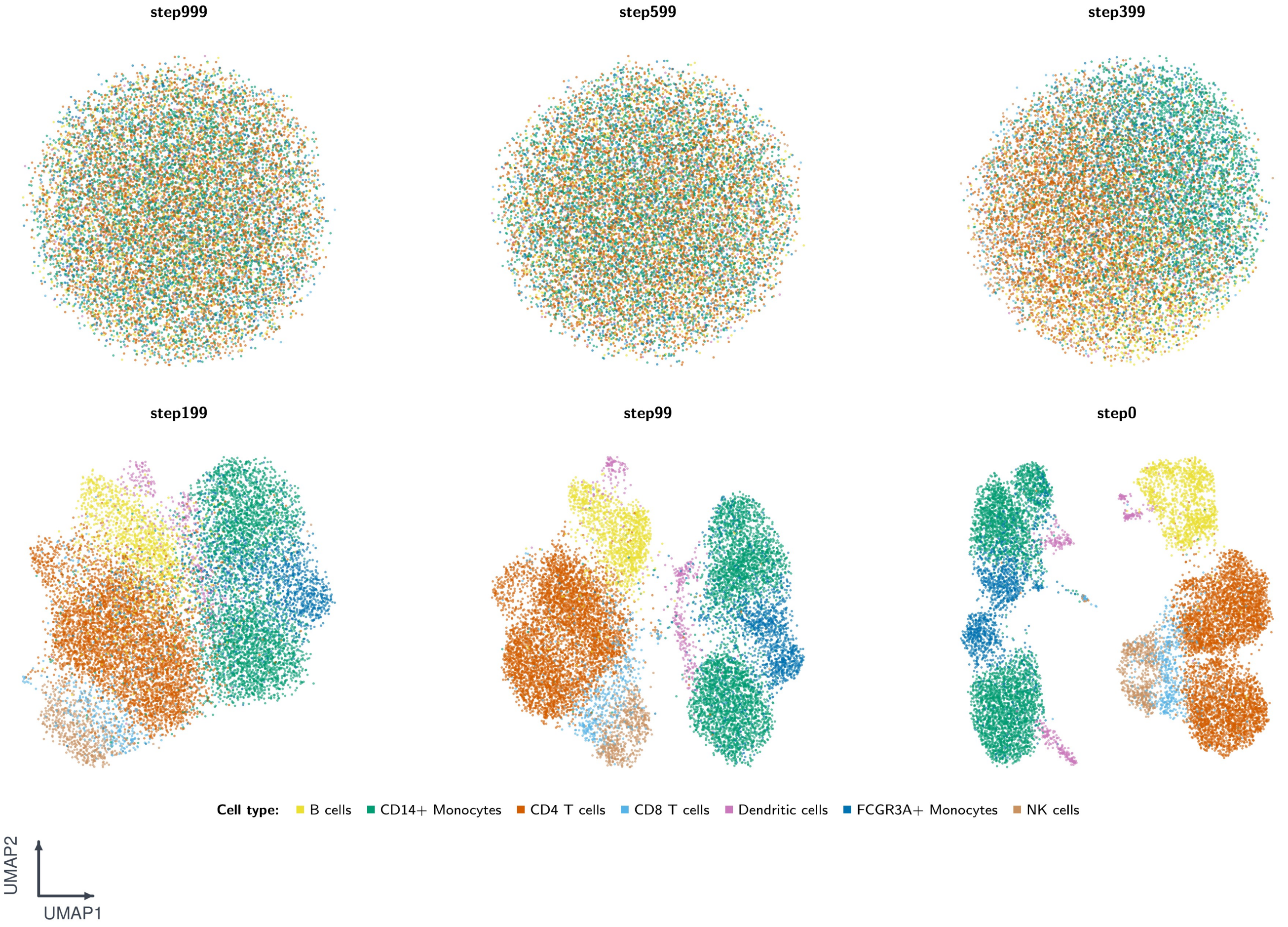


**Supplementary Fig. S11. PBMC reverse-diffusion latent states coloured by cell type.** UMAP visualisations show selected PerturbLDM latent states for the Fig. 5 PBMC IFN-$\beta$ response-generation analysis at step999, step599, step399, step199, step99 and step0. Panel labels report the post-update diffusion index, so the saved latent_1000 snapshot appears as step999. Points are coloured by cell type, with a small UMAP1/UMAP2 indicator for orientation. These embeddings visualise reverse denoising and do not trace a biological trajectory.

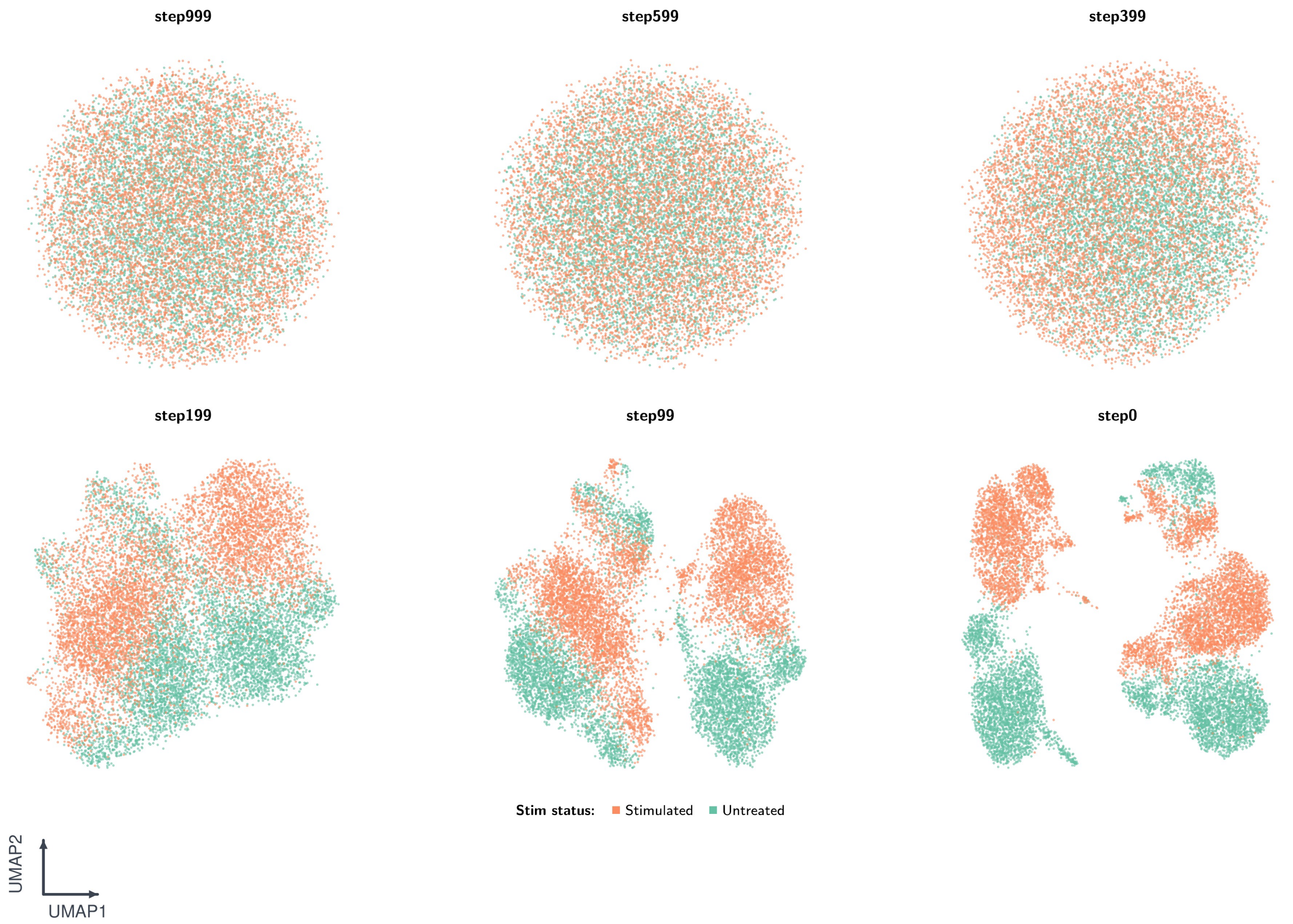


**Supplementary Fig. S12. PBMC reverse-diffusion latent states coloured by stimulation state.** The same selected latent-state snapshots as in Supplementary Fig. S11 are coloured by stimulation annotation and are ordered left-to-right, top-to-bottom as step999, step599, step399, step199, step99 and step0. The UMAP1/UMAP2 indicator denotes embedding orientation. This annotation-specific view is qualitative and does not quantify latent-state trajectories or distributions.

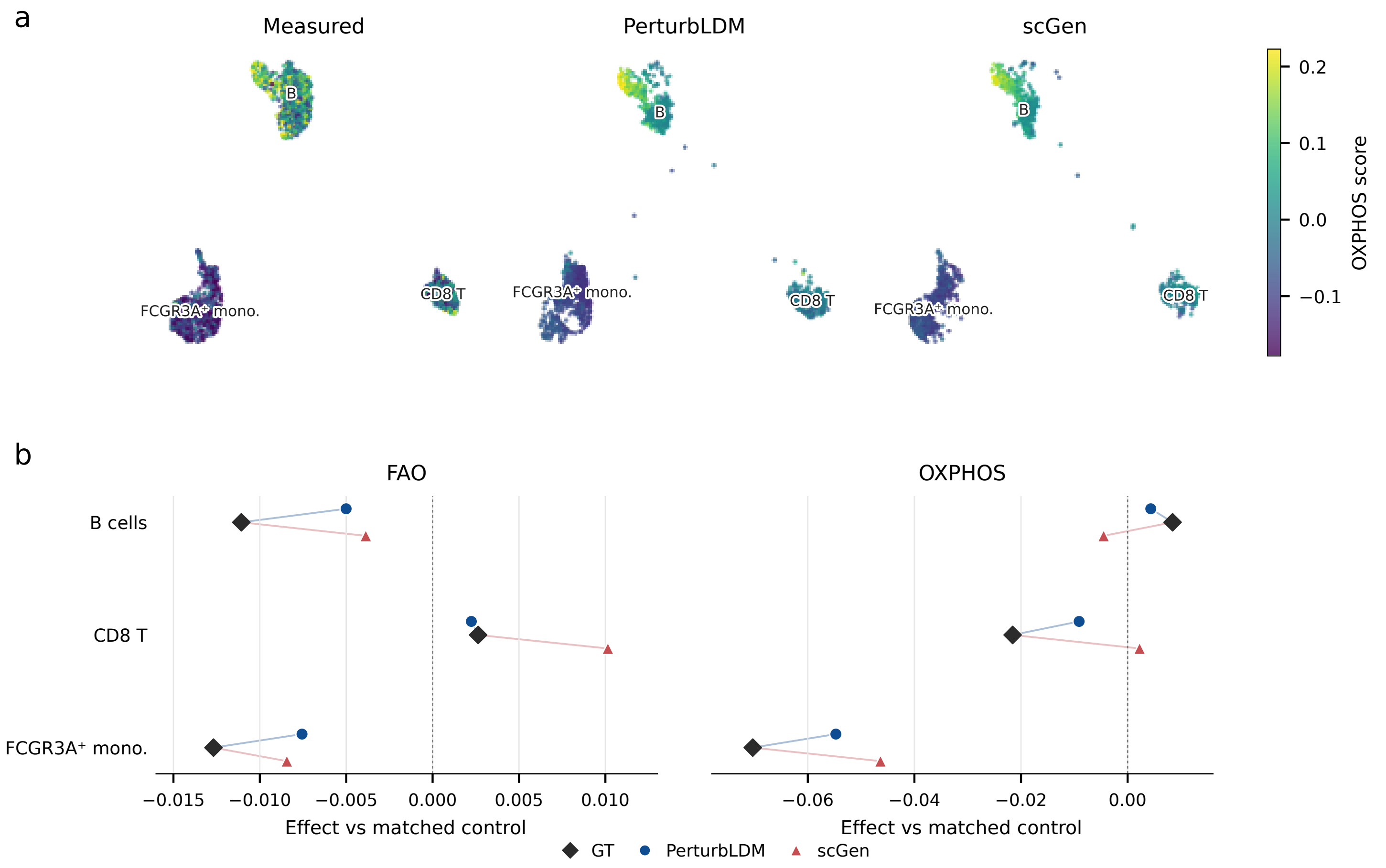


**Supplementary Fig. S13. PBMC FAO and OXPHOS transcriptional programme scores. a,** OXPHOS scores on a measured-reference UMAP. PCA and UMAP were fitted to measured stimulated profiles, and PerturbLDM and scGen profiles were projected through the same models. Panels use identical coordinates and a common colour scale, and labels mark lineage centroids. **b,** Matched-control-relative effects for GO Biological Process fatty-acid beta-oxidation (FAO) and Hallmark OXPHOS across the three held-out lineages. Black diamonds denote measured stimulated profiles, blue circles PerturbLDM and red triangles scGen. Points are vertically offset within each lineage for visibility. Panel **b** shows the programme components underlying the composite effect in Fig. 5h. Scores were computed over 7 and 31 genes, respectively, represented in the shared 2,000-HVG space, and reflect programme-associated gene expression rather than metabolic flux.

#### Supplementary Tables

Table S1: **Goserelin-HCT15 metrics for the held-out 0.5 $\mu$M condition.** The condition was evaluated against CPA and chemCPA using two single-cell distribution metrics and three absolute-expression profile metrics. Distribution metrics were computed with 500 observed and 500 predicted cells. These cells are nested observations within one selected condition, not independent biological replicates. Lower MMD-RBF, OT Wasserstein and MAE values and higher $R^2$ and Pearson $r$ values indicate smaller discrepancies from the observed profile and empirical cell distribution. Bold indicates the best value among methods and does not imply a method-comparison $P$ value.

| | Distribution alignment | | Absolute-expression profile | | |
|---|---|---|---|---|---|
| Method | MMD-RBF ↓ | OT Wasserstein ↓ | $R^2$ ↑ | Pearson $r$ ↑ | MAE ↓ |
| PerturbLDM | **0.2106** | **62.14** | **0.9844** | **0.9923** | **0.0194** |
| chemCPA | 0.2150 | 62.42 | 0.9561 | 0.9786 | 0.0320 |
| CPA | 0.2396 | 63.24 | 0.8705 | 0.9529 | 0.0520 |

Table S2: **PANACEA query-level MoA-neighbour concordance.** For each of 27 MoA-evaluable PANACEA query drugs, candidate drugs were ranked by matched-control-relative Hallmark GSVA pathway-effect similarity after excluding the query drug. Broad Hub MoA annotations were applied only after ranking. Bold rows indicate top-1 endpoint success, not row-wise statistical significance. A dash indicates that the top-ranked neighbour shared no annotated MoA with the query. The combined endpoint was 14 of 27 (51.9%, Wilson 95% CI, 34.0–69.3%, one-sided exact Poisson-binomial $P = 0.1207$).

| Query drug | Top-1 neighbour | Sim. | Shared MoA with top-1 neighbour | First MoA-sharing rank | MoA-sharing candidates |
|---|---|---|---|---|---|
| *Exact drug identity present in Tahoe training records* | | | | | |
| **Afatinib** | **Dacomitinib** | **0.987** | **EGFR inhibitor** | **1** | **9/26** |
| Cabozantinib | Gefitinib | 0.978 | – | 3 | 9/26 |
| Gefitinib | Cabozantinib | 0.978 | – | 3 | 9/26 |
| Lapatinib | Nilotinib | 0.975 | – | 5 | 9/26 |
| **Neratinib** | **Osimertinib** | **0.985** | **EGFR inhibitor** | **1** | **9/26** |
| **Osimertinib** | **Neratinib** | **0.985** | **EGFR inhibitor** | **1** | **9/26** |
| Ponatinib | Gefitinib | 0.975 | – | 4 | 13/26 |
| **Regorafenib** | **Sorafenib** | **0.985** | **KIT inhibitor; PDGFR tyrosine kinase receptor inhibitor; RAF inhibitor; RET tyrosine kinase inhibitor; VEGFR inhibitor** | **1** | **13/26** |
| **Sunitinib** | **Cediranib** | **0.976** | **KIT inhibitor; VEGFR inhibitor** | **1** | **15/26** |
| **Vandetanib** | **Neratinib** | **0.972** | **EGFR inhibitor** | **1** | **16/26** |
| *Exact drug identity absent from Tahoe training records* | | | | | |
| **AEE788-like EGFR/ VEGFR inhibitor** | **Gefitinib** | **0.973** | **EGFR inhibitor** | **1** | **16/26** |
| **Bafetinib** | **Nilotinib** | **0.975** | **BCR-ABL kinase inhibitor** | **1** | **5/26** |
| Bosutinib | Osimertinib | 0.984 | – | 10 | 6/26 |
| **Cediranib** | **Sunitinib** | **0.976** | **KIT inhibitor; VEGFR inhibitor** | **1** | **11/26** |
| **Crenolanib** | **Dasatinib** | **0.984** | **PDGFR tyrosine kinase receptor inhibitor** | **1** | **8/26** |
| **Dacomitinib** | **Afatinib** | **0.987** | **EGFR inhibitor** | **1** | **9/26** |
| **Dasatinib** | **Crenolanib** | **0.984** | **PDGFR tyrosine kinase receptor inhibitor** | **1** | **13/26** |
| **Dovitinib** | **Neratinib** | **0.969** | **EGFR inhibitor** | **1** | **22/26** |
| Foretinib | Bosutinib | 0.953 | – | 4 | 9/26 |
| Icotinib | Linifanib | 0.950 | – | 2 | 9/26 |
| Imatinib | KW-2449 | 0.951 | – | 3 | 12/26 |
| KW-2449 | Imatinib | 0.951 | – | 4 | 7/26 |
| Linifanib | Lapatinib | 0.970 | – | 6 | 13/26 |
| Nilotinib | Lapatinib | 0.975 | – | 2 | 6/26 |
| Quizartinib | Nilotinib | 0.943 | – | 9 | 5/26 |
| **Sorafenib** | **Regorafenib** | **0.985** | **KIT inhibitor; PDGFR tyrosine kinase receptor inhibitor; RAF inhibitor; RET tyrosine kinase inhibitor; VEGFR inhibitor** | **1** | **15/26** |
| Tivantinib | Bafetinib | 0.954 | – | 26 | 1/26 |

**Table S3: Selected PBMC IFN-$\beta$ GO Biological Process terms and gene membership.** The seven terms were selected from the measured-response analysis for the focused display. Official GO term names are from QuickGO and gene membership is from the Enrichr GO Biological Process 2023 library. Membership documents the annotation set and is not evidence of model recovery.

| GO term and source | Gene symbols |
|---|---|
| **defense response to virus**<br>GO:0051607<br>`http://amigo.geneontology.org/amigo/term/GO:0051607`<br>189 genes | ACOD1, AGBL4, AGBL5, AICDA, AIM2, AKAP1, APOBEC3A, APOBEC3B, APOBEC3C, APOBEC3D, APOBEC3F, APOBEC3G, APOBEC3H, ATAD3A, ATG7, AZI2, AZU1, BCL2, BCL2L1, BNIP3, BNIP3L, BPIFA1, BST2, CARD8, CASP1, CD207, CD40, CGAS, CHMP3, CLPB, CNOT7, CXCL10, DDIT4, DDX56, DDX60, DDX60L, DHX15, DHX16, DHX58, DUS2, EIF2AK2, EXOSC4, EXOSC5, F2RL1, FADD, FCN3, G3BP1, GARIN5A, GBP2, GBP5, GBP7, IFI16, IFI27, IFI44L, IFI6, IFIH1, IFIT1, IFIT1B, IFIT2, IFIT3, IFIT5, IFITM1, IFITM2, IFITM3, IFNA2, IFNAR2, IFNB1, IFNE, IFNL1, IFNL2, IFNL3, IFNL4, IFNLR1, IGF2BP1, IKBKE, IL21, IL6, IRF1, IRF2, IRF3, IRF7, ISG15, ISG20, LILRB1, LYST, MAP3K14, MARCHF2, MAVS, MBL2, MICA, MID2, MLKL, MORC3, MOV10, MX1, MX2, MYD88, NCBP1, NCBP3, NCK1, NDUFAF4, NLRC5, NLRP1, NLRP6, NLRP9, NMB, NMBR, NT5C3A, OAS1, OAS2, OAS3, OASL, OPRK1, PDE12, PHB1, PHB2, PLA2G10, PLSCR1, PMAIP1, PML, PQBP1, PRF1, PTPRC, PYCARD, RAB2B, RELA, RIGI, RIPK3, RNASE1, RNASE2, RNASE6, RNASEL, RNF135, RNF185, RSAD2, RTP4, SAMHD1, SENP7, SERINC3, SERINC5, SETD2, SHFL, SKP2, SLFN11, SLFN13, STAT1, STAT2, STING1, TANK, TBK1, TBKBP1, TLR2, TLR3, TLR7, TLR8, TLR9, TNF, TRAF3, TRAF6, TRIM11, TRIM13, TRIM15, TRIM21, TRIM25, TRIM26, TRIM28, TRIM31, TRIM32, TRIM35, TRIM41, TRIM5, TRIM52, TRIM56, TRIM6, TRIM7, TRIM8, TTC4, UBE2N, UBE2W, USP18, USP20, USP27X, USP29, USP44, ZCCHC3, ZDHHC1, ZDHHC11, ZMYND11, ZNFX1 |
| **negative regulation of viral genome replication**<br>GO:0045071<br>`http://amigo.geneontology.org/amigo/term/GO:0045071`<br>53 genes | APOBEC3A, APOBEC3B, APOBEC3C, APOBEC3D, APOBEC3F, APOBEC3G, APOBEC3H, BANF1, BST2, BTBD17, CCL5, EIF2AK2, FAM111A, HMGA2, IFI16, IFIH1, IFIT1, IFIT5, IFITM1, IFITM2, IFITM3, IFNB1, IFNL3, ILF3, INPP5K, ISG15, ISG20, LTF, MAVS, MORC2, MPHOSPH8, MX1, N4BP1, OAS1, OAS2, OAS3, OASL, PLSCR1, PROX1, RESF1, RNASEL, RSAD2, SETDB1, SHFL, SLPI, SRPK1, SRPK2, TASOR, TNF, TNIP1, TRIM6, ZC3HAV1, ZNFX1 |
| **response to type II interferon**<br>GO:0034341<br>`http://amigo.geneontology.org/amigo/term/GO:0034341`<br>80 genes | AIF1, AQP4, BST2, CALCOCO2, CALM1, CAMK2A, CASP1, CCL1, CCL11, CCL13, CCL14, CCL15, CCL16, CCL17, CCL18, CCL19, CCL2, CCL20, CCL21, CCL22, CCL23, CCL24, CCL25, CCL26, CCL3, CCL3L1, CCL4, CCL4L1, CCL5, CCL7, CCL8, CD40, CD47, CD58, CD74, CIITA, CITED1, CX3CL1, CXCL16, CYP27B1, DAPK1, DAPK3, EPRS1, GAPDH, GBP1, GBP2, GBP4, GBP5, GBP6, GCH1, HLA-DPA1, IFITM1, IFITM2, IFITM3, IL12RB1, IL23R, IRF8, KYNU, LGALS9, MEFV, NUB1, PDE12, RPL13A, SHFL, SIRPA, SLC11A1, SLC22A5, SLC26A6, SNCA, SP100, STAT1, SYNCRIP, TDGF1, TLR2, TLR4, TRIM21, UBD, WNT5A, XCL1, XCL2 |
| **cellular response to type II interferon**<br>GO:0071346<br>`http://amigo.geneontology.org/amigo/term/GO:0071346`<br>66 genes | AIF1, AQP4, CALM1, CAMK2A, CASP1, CCL1, CCL11, CCL13, CCL14, CCL15, CCL16, CCL17, CCL18, CCL19, CCL2, CCL20, CCL21, CCL22, CCL23, CCL24, CCL25, CCL26, CCL3, CCL3L1, CCL4, CCL4L1, CCL5, CCL7, CCL8, CD47, CD58, CX3CL1, DAPK1, DAPK3, EPRS1, GAPDH, GBP1, GBP2, GBP4, GBP5, GBP6, HCK, HLA-DPA1, IFNG, IFNGR1, IFNGR2, IL12RB1, IRF1, IRF8, JAK1, JAK2, LGALS9, PDE12, RPL13A, SIRPA, SLC26A6, SP100, STAT1, SYNCRIP, TDGF1, TLR2, TLR4, TYK2, WNT5A, XCL1, XCL2 |
| **negative regulation of viral process**<br>GO:0048525<br>`http://amigo.geneontology.org/amigo/term/GO:0048525`<br>61 genes | APOBEC3A, APOBEC3C, APOBEC3F, APOBEC3G, APOBEC3H, BANF1, BST2, BTBD17, CCL5, EIF2AK2, FAM111A, HEXIM1, IFI16, IFIH1, IFIT1, IFIT5, IFITM1, IFITM2, IFITM3, IFNB1, IFNL3, ILF3, ISG15, ISG20, LARP7, LTF, MAVS, MBL2, MID2, MX1, N4BP1, OAS1, OAS2, OAS3, OASL, PITX3, PLSCR1, PPIA, PROX1, RNASEL, RSAD2, SHFL, SLPI, SRPK1, SRPK2, STAT1, TNF, TNIP1, TRIM11, TRIM13, TRIM14, TRIM21, TRIM27, TRIM31, TRIM32, TRIM6, TRIM62, UBP1, ZC3HAV1, ZFP36, ZNFX1 |

Continued on next page

Table S3 (continued)

| GO term and source | Gene symbols |
|---|---|
| **regulation of viral genome replication**<br>GO:0045069<br>http://amigo.geneontology.org/amigo/term/GO:0045069<br>67 genes | ADARB1, APOBEC3A, APOBEC3C, APOBEC3F, APOBEC3G, BANF1, BST2, BTBD17, CCL5, CD28, CNOT7, CXCL8, DDB1, DDX3X, EIF2AK2, FAM111A, FKBP6, GBP7, HACD3, IFI16, IFIH1, IFIT1, IFIT5, IFITM1, IFITM2, IFITM3, IFNB1, IFNL3, ILF3, ISG15, ISG20, LARP1, LTF, MAVS, MX1, N4BP1, NR5A2, OAS1, OAS2, OAS3, OASL, PABPC1, PDE12, PKN2, PLSCR1, PPARA, PPIA, PPID, PPIE, PPIH, PROX1, RAD23A, RNASEL, RSAD2, SHFL, SLPI, SRPK1, SRPK2, STAU1, TARBP2, TMEM39A, TNF, TNIP1, TRIM6, VAPB, ZC3HAV1, ZNFX1 |
| **response to type I interferon**<br>GO:0034340<br>http://amigo.geneontology.org/amigo/term/GO:0034340<br>9 genes | IFIT1, ISG15, MX1, SETD2, SHFL, SHMT2, SMPD1, SP100, TRIM56 |

Table S4: **Condition-paired Tahoe comparisons T01–T09.** Complete direction-oriented effect sizes, uncertainty intervals and multiplicity-adjusted tests underlying Main Fig. 2 are reported here. Each comparison contains the same 13,942 held-out drug-dose-cell-line conditions. Positive improvements favour PerturbLDM. Confidence intervals are two-sided percentile 95% intervals for the median paired improvement from 2,000 condition bootstrap resamples (seed 20260718). $P$ values are from two-sided paired Wilcoxon signed-rank tests with Benjamini–Hochberg adjustment within the three effect, two signed-Hallmark and four distribution comparisons. These statistics describe variation across the held-out condition set and do not estimate variation across model retraining or independent atlases. Analysis units and denominators are summarised in Table S5.

| Family | Comparator | Endpoint | Median improvement (95% CI) | Favoured (%) | BH-adjusted $P$ |
|---|---|---|---|---|---|
| Matched-control effect | AdditiveMean | all-gene effect Pearson ↑ | 0.08085 (0.07957–0.08215) | 95.23 | $< 10^{-300}$ |
| Matched-control effect | AdditiveMean | all-gene effect Spearman ↑ | 0.07903 (0.07804–0.08000) | 98.01 | $< 10^{-300}$ |
| Matched-control effect | AdditiveMean | effect MAE ↓ | 0.004670 (0.004617–0.004725) | 98.98 | $< 10^{-300}$ |
| Signed Hallmark effect | AdditiveMean | profile Pearson ↑ | 0.1162 (0.1143–0.1187) | 94.36 | $< 10^{-300}$ |
| Signed Hallmark effect | AdditiveMean | score MAE ↓ | 0.003640 (0.003555–0.003707) | 87.05 | $< 10^{-300}$ |
| Distribution fidelity | CPA | MMD-RBF ↓ | 0.03489 (0.03477–0.03499) | 100.0 | $< 10^{-300}$ |
| Distribution fidelity | chemCPA | MMD-RBF ↓ | 0.01138 (0.01128–0.01146) | 99.98 | $< 10^{-300}$ |
| Distribution fidelity | CPA | OT Wasserstein ↓ | 1.357 (1.353–1.361) | 100.0 | $< 10^{-300}$ |
| Distribution fidelity | chemCPA | OT Wasserstein ↓ | 0.5010 (0.4965–0.5059) | 99.96 | $< 10^{-300}$ |

Table S5: **Analysis design and evaluation units.** This table summarises the unit, denominator, data definition and interpretation for each analysis module.

| Analysis | Unit and denominator | Design and data definition | Interpretation |
|---|---|---|---|
| Tahoe benchmark split | 32,529 development and 13,942 held-out evaluation conditions | Each condition is one drug-dose-cell-line combination. All 379 drugs, 47 cell lines and three dose levels occur in both splits. | Benchmark of unmeasured combinations of observed factors. |
| Learned-comparator selection | 29,277 fitting and 3,252 validation conditions; 13,942 held-out evaluation conditions | MLP, random forest, CPA and chemCPA use the same condition membership. Validation selects hyperparameters or model epochs; the held-out evaluation set is excluded from selection. | Final comparison uses the selected setting for each method. |
| Matched-control effects (T01–T03) | 13,942 paired held-out conditions | All-gene effect Pearson correlation, Spearman correlation and MAE are compared with AdditiveMean. | The held-out condition is the paired analysis unit. |
| Signed Hallmark effects (T04–T05) | 13,942 paired held-out conditions; 50 Hallmark gene sets | Signed mean matched-control effects are computed over the members of each Hallmark set. | Programme-level concordance and error; not an enrichment test. |
| Cell-state distributions (T06–T09) | 13,942 paired conditions per method; 500 observed and 500 predicted cells per condition | MMD-RBF and OT Wasserstein distances compare condition-aligned empirical cell samples. | Local distributional concordance, not calibration of the full conditional distribution. |
| PANACEA | 64 grouped response profiles; 27 MoA-evaluable query drugs | Profiles are aligned with matched controls, converted to Hallmark pathway effects and ranked after same-drug exclusion. | Descriptive pharmacological-neighbourhood analysis. |
| Fetal colon | One pooled broad-stage-2 comparison; 1,529 held-out cells; 800 genes | Early and mature enterocyte states provide the observed anchors; both methods use the same gene set. | Study-specific developmental-state comparison with descriptive PCW stratification. |
| PBMC expression and DEG | Three held-out stimulated lineages with retained unstimulated controls | The same 2,000-gene space and held-out stimulated profiles are used for PerturbLDM, PerturbLDM-ctrl and scGen. | Descriptive lineage-level comparison ($n = 3$ lineages). |
| PBMC focused GO programmes | Seven author-selected terms from the measured-response analysis | GO Biological Process 2023 membership is applied consistently across measured and predicted profiles. | Focused descriptive programme recovery. |

**Table S6: Detailed summary values for the fetal-colon and PBMC applications.** Gene- and lineage-level descriptive values corresponding to Main Figs. 4 and 5 are reported. Fetal-colon intervals are gene-level bootstrap summaries over correlated genes and do not provide biological-replicate inference. PBMC values are shown by lineage before any unweighted mean. PerturbLDM-ctrl is a sensitivity analysis using strength 0.4 selected against the held-out targets.

#### a. Fetal-colon gene-level descriptive summaries

| Endpoint | PerturbLDM | Squidiff | Direction-oriented difference (95% gene-bootstrap interval) | Analysis unit and scope |
|---|---|---|---|---|
| Pseudo-bulk absolute mean-expression error, median | 0.01069 | 0.07743 | 0.06398 (0.06118–0.06746) reduction | 800 genes; descriptive |
| Single-gene Wasserstein distance, median | 0.08137 | 0.1300 | 0.04576 (0.04322–0.04884) reduction | 800 genes; descriptive |
| PCW-stratified mean-expression MAE, median | 0.02003 | 0.08878 | 0.05400 (0.05057–0.05709) reduction | 800 genes; descriptive PCW strata |
| Fraction with absolute error $\leq 0.1$ | 0.9313 (745/800) | 0.6838 (547/800) | 0.2475 increase; no interval reported | gene counts, not biological replicates |

##### b. PBMC stimulated-state expression metrics by held-out lineage

| Lineage | Method | Pearson ↑ | Spearman ↑ | $R^2$ ↑ | RMSE ↓ | MAE ↓ |
|---|---|---|---|---|---|---|
| B cells | scGen | 0.9675 | 0.8368 | 0.9330 | 0.0992 | 0.0502 |
| B cells | PerturbLDM | 0.9731 | 0.9007 | 0.9451 | 0.0898 | 0.0417 |
| B cells | PerturbLDM-ctrl (sensitivity) | 0.9712 | 0.9002 | 0.9417 | 0.0926 | 0.0418 |
| CD8 T cells | scGen | 0.9730 | 0.8471 | 0.9454 | 0.0861 | 0.0464 |
| CD8 T cells | PerturbLDM | 0.9841 | 0.9032 | 0.9649 | 0.0690 | 0.0350 |
| CD8 T cells | PerturbLDM-ctrl (sensitivity) | 0.9857 | 0.9047 | 0.9692 | 0.0646 | 0.0339 |
| FCGR3A$^+$ monocytes | scGen | 0.9789 | 0.8275 | 0.9502 | 0.1235 | 0.0492 |
| FCGR3A$^+$ monocytes | PerturbLDM | 0.9840 | 0.9213 | 0.9679 | 0.0991 | 0.0334 |
| FCGR3A$^+$ monocytes | PerturbLDM-ctrl (sensitivity) | 0.9852 | 0.9213 | 0.9699 | 0.0961 | 0.0331 |

##### c. PBMC matched-control DEG-set metrics by held-out lineage

| Lineage | Method | Recall (%) | Precision (%) | Specificity (%) | F1 (%) |
|---|---|---|---|---|---|
| B cells | scGen | 82.4 | 32.6 | 43.6 | 46.7 |
| B cells | PerturbLDM | 81.5 | 46.3 | 68.7 | 59.1 |
| B cells | PerturbLDM-ctrl (sensitivity) | 81.5 | 44.9 | 66.9 | 57.9 |
| CD8 T cells | scGen | 88.2 | 12.9 | 44.8 | 22.5 |
| CD8 T cells | PerturbLDM | 88.2 | 12.4 | 42.7 | 21.8 |
| CD8 T cells | PerturbLDM-ctrl (sensitivity) | 88.2 | 14.2 | 50.5 | 24.4 |
| FCGR3A$^+$ monocytes | scGen | 81.6 | 63.5 | 46.2 | 71.4 |
| FCGR3A$^+$ monocytes | PerturbLDM | 90.3 | 60.8 | 33.3 | 72.7 |
| FCGR3A$^+$ monocytes | PerturbLDM-ctrl (sensitivity) | 88.8 | 63.7 | 42.1 | 74.2 |
